# RNA partitioning into stress granules is based on the summation of multiple interactions

**DOI:** 10.1101/2020.04.15.043646

**Authors:** Tyler Matheny, Briana Van Treeck, Thao Ngoc Huynh, Roy Parker

**Affiliations:** Department of Biochemistry, University of Colorado, Boulder, CO 80309, USA; Howard Hughes Medical Institute, Chevy Chase, MD 20815, USA

**Author notes:** These authors contributed equally.

## Abstract

Stress granules (SGs) are stress-induced RNA-protein assemblies formed from a complex transcriptome of untranslating ribonucleoproteins (RNPs). Although RNAs can be either enriched or depleted from SGs, the rules that dictate RNA partitioning into SGs are unknown. We demonstrate that the SG-enriched *NORAD* RNA is sufficient to enrich a reporter RNA within SGs through the combined effects of multiple elements. Moreover, artificial tethering of G3BP1 or TIA1 can target mRNAs into SGs in a dose-dependent manner that suggests individual protein interactions have small effects on the SG partitioning of mRNPs, which is supported by the observation that the SG transcriptome is largely unchanged in cell lines lacking the abundant SG RNA-binding proteins G3BP1 and G3BP2. We suggest the targeting of RNPs into SGs is due to a summation of potential RNA-protein, protein-protein, and RNA-RNA interactions with no single interaction dominating RNP recruitment into SGs.

## INTRODUCTION

Stress granules (SGs) are ribonucleoprotein (RNP) assemblies that form during stress when translation initiation is limited (Panas et al., 2016; Protter and Parker, 2016). SGs are of interest because they play roles in the stress response, are related to similar RNP granules in neurons and embryos, share components with toxic aggregates observed in degenerative disease, and affect viral infections as well as cancer progression (Anderson et al., 2015; Kedersha et al., 2013; Kim et al., 2013; Li et al., 2013; Reineke and Lloyd, 2013; Somasekharan et al., 2015).

Based on super-resolution microscopy, mammalian SGs formed during arsenite stress are non-uniform in nature and contain local regions of protein and RNA concentration, referred to as SG cores (Jain et al., 2016; Niewidok et al., 2018; Wheeler et al., 2016). SG cores are stable sub-assemblies in cell lysates (Jain et al., 2016; Wheeler et al., 2016) allowing for their purification from U-2 OS cells expressing GFP-G3BP through differential centrifugation and GFP pulldown (Wheeler et al., 2017; Khong et al., 2018). SG purification led to the discovery of a diverse SG proteome composed of numerous RNA-binding proteins (RBPs) forming a dense protein-protein interaction network (Jain et al., 2016). Similar SG protein composition was also detected by proximity labeling methods (Youn et al., 2018; Markmiller et al., 2018). The fact that SGs are composed of a dense protein-protein interaction network is consistent with the recruitment of proteins to SGs through protein-protein interactions, which might recruit specific mRNAs bound to those RBPs.

An additional possibility is that RNA-RNA interactions contribute to defining the RNA composition of SGs. Consistent with this model, the transcriptome of protein-free *in vitro* RNA assemblies, formed from total yeast RNA, shows remarkable overlap with the transcriptome of yeast SGs (Van Treeck et al., 2018). Moreover, the abundant RNA helicase eIF4A functions to limit RNA condensation and SG formation by binding RNA (Tauber et al., 2020). This suggests a model wherein RNPs are partitioned into SGs by both RNA-RNA and protein-protein interactions.

A striking feature of the SG transcriptome is that there are dramatic differences in the partitioning of RNPs into SGs. For example, RNA sequencing of immunopurified SGs revealed that some RNAs, such as *AHNAK* or *NORAD*, are strongly enriched in SGs, while other mRNAs, such as *GAPDH*, are largely depleted from SGs (Khong et al., 2017). Differential partitioning of RNPs into SGs and/or P-bodies has also been seen when RNP granules are fractioned based on particle sorting or differential centrifugation (Namkoong et al., 2018; Matheny et al., 2019; Hubstenberger et al., 2017). These analyses revealed that long, poorly translated transcripts preferentially enrich in SGs (Khong et al., 2017), that AU-rich elements correlated with SG granule fractionation (Namkoong et al., 2018), and that decreased translational efficiency correlated with increased SG and P-body enrichment (Khong et al., 2017; Hubstenberger et al., 2017; Matheny et al., 2019). Taken together, these observations suggest that partitioning of RNPs into SGs could be promoted by protein- or RNA-based interactions, affected by overall length of the RNA, and inhibited by the association with ribosomes. However, the relative importance of these interactions, their required valency, and how they might contribute to the RNA composition of, and organization within, SGs is unknown.

Herein, we examine the rules that affect the partitioning of RNAs into SGs. We first show enrichment into SGs is a dominant property as a chimera between the highly enriched *NORAD* RNA and a poorly localized reporter RNA is enriched in SGs. Deletion analysis revealed the enrichment of *NORAD* in SGs is due to multiple sequence elements that act in an additive manner. In addition, we examined how proteins influence the recruitment of RNAs to SGs by determining how tethering of RBPs found in SGs to a reporter transcript affects mRNA enrichment in SGs and how deletion of the major SG RBPs G3BP1 and G3BP2 affects RNA recruitment into SGs. We show that artificially tethering multiple G3BP1 molecules to a *luciferase* reporter transcript increases the SG localization of a *luciferase* RNA reporter in a dose-dependent manner. However, based on curve fitting of the G3BP1 tethering experiments, we predicted that removal of G3BP proteins from the cell would only have a limited effect on the RNA composition of SGs. Testing this hypothesis, we demonstrate that the RNA composition of sorbitol-induced SGs is very similar in wild type (WT) and ΔΔG3BP1/2 cell lines. Taken together, our results indicate that while G3BP1 binding can promote RNA targeting to SGs, it is not required for SG localization. We suggest a model in which RNA localization to SGs arises through the summation of many RNA-RBP and RNA-RNA interactions, which each individually play a small role in localization, but through synergistic effects lead to the observed RNP partitioning into RNP granules.

## RESULTS

### *Luciferase* mRNA as a reporter for SG partitioning in mammalian cells

The firefly *luciferase* was used as an RNA reporter to assess how different elements affect RNP partitioning into SGs for two reasons. First, *luciferase* is not endogenous in mammalian cells, thus, we can eliminate the endogenous background in our analysis. Second, single molecule fluorescent *in situ* hybridization (smFISH) analyses revealed that *luciferase* mRNA is poorly enriched in SGs, with approximately 15% of its cytoplasmic RNA molecules in SGs upon an hour of sodium arsenite treatment (**Figure S1**). Using this reporter system, we sought to test how adding sequences or binding sites for RBPs to this reporter affected SG partitioning.

### *NORAD* RNA contains dominant elements that dictate SG partitioning

While it is assumed that partitioning of RNPs into SGs is a dominant trait, to our knowledge this has never been tested. To determine if SG partitioning is a dominant trait we inserted the SG-enriched *NORAD* RNA into the 3’ UTR of the *luciferase* reporter mRNA. As assessed by smFISH, we observed that ∼71% of the chimeric *luciferase-NORAD* RNA was recruited to SGs, similar to the endogenous *NORAD* RNA (76.7%) and significantly increased compared to the *luciferase* reporter (16.7%) (**Figure 1A-C**). Thus, SG recruitment of RNPs is a dominant property.

**Figure 1:**
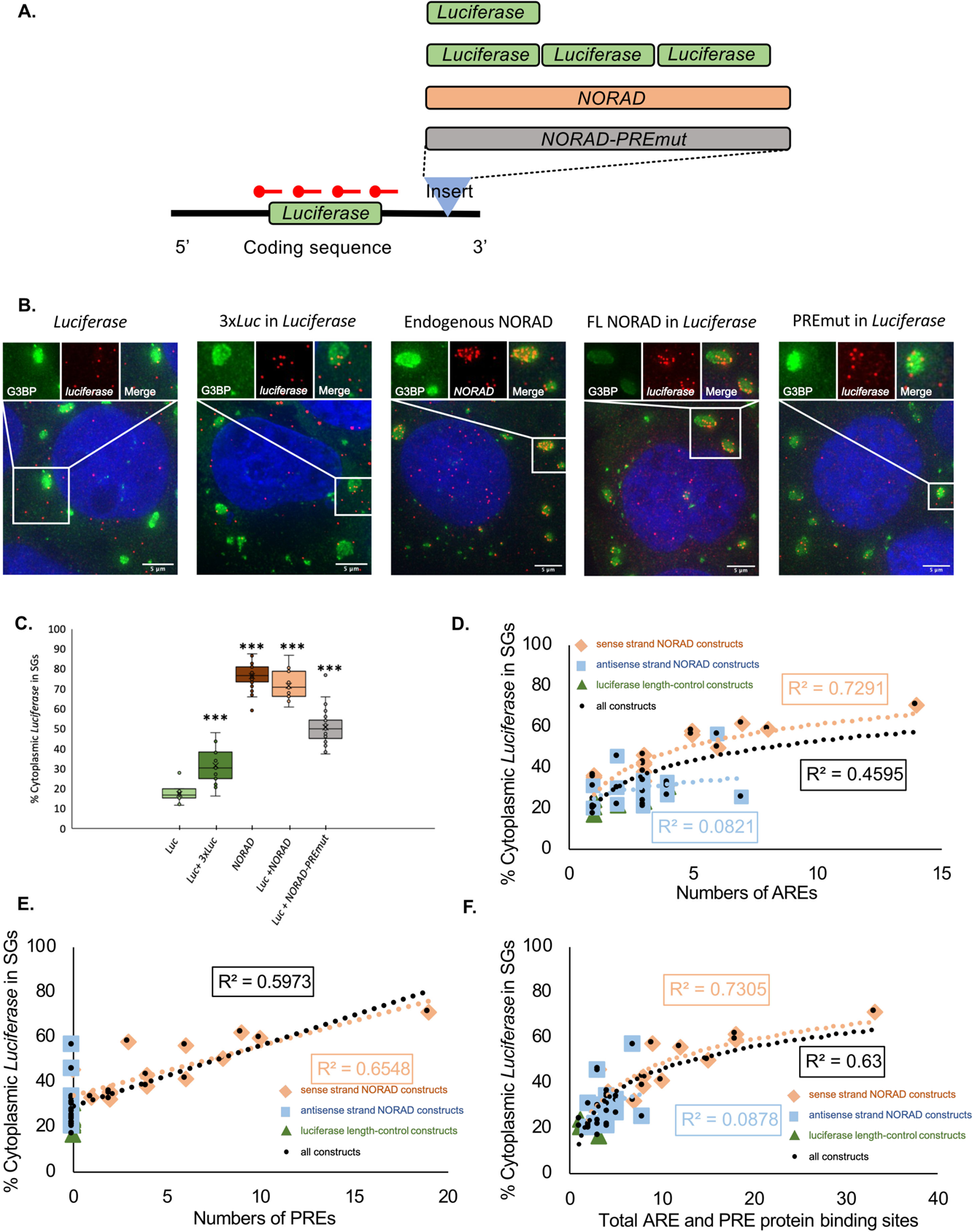
*NORAD* increases *luciferase* RNA enrichment within SGs **(A)** Cartoon depicting chimeric *luciferase* reporter constructs with different inserts in the 3’ UTR. PREmut is the *NORAD* sequence with 18 of the PREs mutated. The 3x luciferase was inserted as an approximate length control for NORAD. **(B)** smFISH of *luciferase* RNA for different constructs during arsenite stress. The endogenous *NORAD* RNA was imaged with smFISH probes to *NORAD*, all other constructs were imaged with smFISH probes to *luciferase*. Scale bar = 5 µm. **(C)** Boxplot of the enrichment of luciferase with different 3’ UTR inserts and endogenous NORAD in SGs. ** p<1×10^-4^. **(D)** Correlation between SG enrichment and number of AREs. (**E**) Correlation between SG enrichment and number of PREs. (**F**) Correlation between SG enrichment and total number of AREs and PREs. For (**D**-**F**): Orange diamonds are sense-NORAD constructs, light blue squares are antisense-NORAD constructs, dark green triangles are luciferase length control constructs, and black circles represent the constructs.

To determine whether one or more elements in the *NORAD* RNA were promoting SG partitioning, we constructed *luciferase* mRNAs with either each half, each of the four quarters, or each eighth of the *NORAD* RNA inserted into the same site in the 3’ UTR.

We observed that essentially any piece of *NORAD* could increase the recruitment of the *luciferase* reporter into SGs with the average increase correlating with the size of the insert (**Supplemental Table S1**). For example, either half of NORAD yielded ∼60% reporter enrichment within SGs, each quarter gave between ∼55% and 35% recruitment, and each eighth gave between 22 and 50% recruitment (**Supplemental Table S1**). These increases are not solely due to increased length since insertion of additional luciferase sequences of similar lengths in the same position within the 3’ UTR gave only limited increases in SG recruitment (**Supplemental Table S1**). Similarly, insertion of antisense sequences from *NORAD* typically resulted in less recruitment of the reporter into SGs than corresponding sense-strand inserts (**Supplemental Table S1**). The limited correlation with length is consistent with the SG transcriptome in which there is an overall length determinant, however mRNAs of similar length can show differences in their partitioning into SGs (Khong et al., 2017). This suggests that *NORAD* contains multiple sequence-specific elements, independent of overall length, that can increase partitioning into SGs.

We compared the features of the *NORAD-luciferase* chimeric RNAs to determine if any feature strongly correlated with SG enrichment. As expected, we observed only a limited correlation between SG recruitment and length or GC content (**Figure S2**). In the *NORAD* RNA there are 13 AU-rich elements (AREs) with an AUUUA motif and 19 Pumilio recognition elements (PREs), which can bind Pumilio and other RBPs (Lee et al., 2016, Tichon et al., 2016). We observed some correlation of SG enrichment in each construct with either the numbers of AREs, PREs, or their summation (**Figure 1D-F**). This implied that elements promoting SG enrichment within *NORAD* might be working by recruiting RBPs.

To test whether the 19 PREs are important for *NORAD* recruitment in SGs, we generated a chimeric mRNA with the *NORAD* sequence with 18 PREs mutated to limit Pumilio (and possibly other proteins) binding, into the *luciferase* 3’ UTR. We observed a significant decrease in *luciferase* reporter, from 71% to 50%, in SGs (**Figure 1B,C**). Since cells lacking Pumilio proteins can still robustly accumulate *NORAD* in SGs (Namkoong et al., 2018), this suggests that the PREs can promote SG enrichment either as an RNA element, or potentially by binding other RBPs, such as SAM68, which has binding sites in or near many of the PREs (Tichon et al., 2018).

Taken together, these observations argue that *NORAD* contains multiple elements that can increase an mRNAs partitioning into SGs. Moreover, the correlation of enrichment with protein binding sites suggests that at least some of the SG enrichment will be due to RBPs targeting mRNAs to SGs.

### Tethering G3BP1 to a reporter mRNA increases the reporter RNA’s enrichment within stress granules

To directly examine how the interactions with mRNA-binding proteins affect mRNA partitioning into SGs, we next focused on G3BP1 for three reasons. First, G3BP1 is one of the most abundant cytoplasmic RBPs (Itzhak et al., 2016; Khong and Parker, 2020). Second, G3BP1 partitions strongly into SGs (Wheeler et al., 2017). Third, G3BP1, and its paralog G3BP2, are required for SG formation under certain stresses (Tourrière et al., 2003; Kedersha et al., 2016). We hypothesized that if protein-RNA interactions of RBPs play a critical role in localizing transcripts to SGs, increasing the number of G3BP1 molecules bound to a specific mRNA by artificial tethering should lead to increased targeting of that mRNA to SGs.

To determine if G3BP1 could increase a given RNA’s localization to SGs, smFISH was used to monitor localization of the *luciferase* RNA reporter with and without G3BP1 tethering sites. We used the λN-BoxB system to artificially tether G3BP1 to the *luciferase* mRNA (Baron-Benhamou et al., 2004). λN was fused to G3BP1-GFP as well as a GFP control (**Figure 2A**). *Luciferase* mRNA reporters were created with 0, 7 and 25 BoxB sequences in their 3’ UTR and were genomically incorporated into the AAVS locus of U-2 OS cells (**Figure 2A**). Reporter RNA localization was assayed in cells transfected with either G3BP1-GFP-λN or GFP-λN and in non-transfected cells.

**Figure 2:**
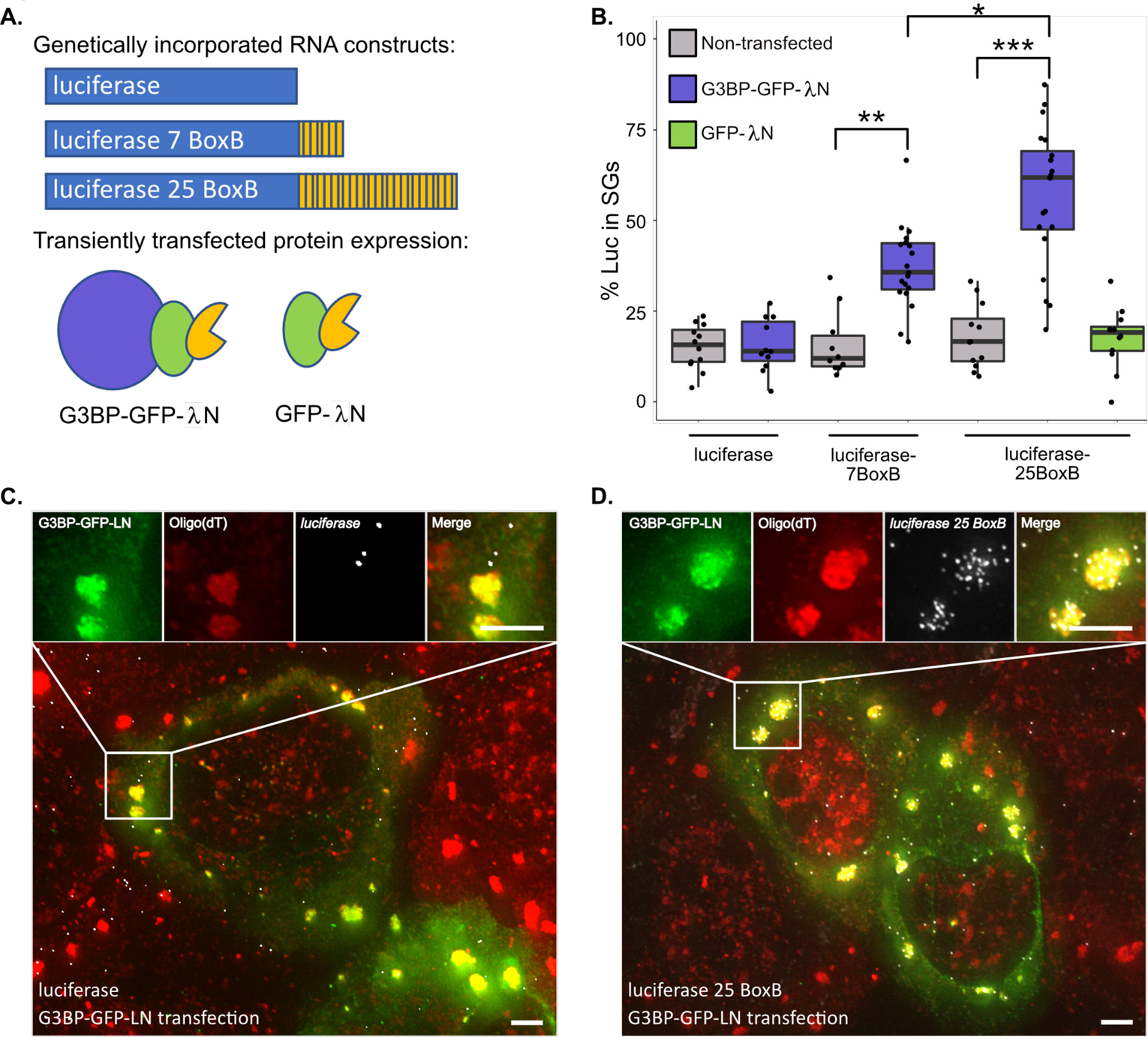
G3BP tethering increases *luciferase* RNA enrichment within SGs **(A)** *Top:* Cartoon depicting genomically incorporated *luciferase* RNA constructs with 0, 7, and 25 BoxB sites in the 3’UTR. *Bottom:* Cartoon depicting G3BP-GFP-λN and GFP-λN used in tethering experiments. **(B)** Boxplot of the enrichment of *luciferase* 0 BoxB, 7 BoxB, and 25 BoxB RNA in non-transfected, G3BP-GFP-λN and GFP-λN transfected cells. * p<0.001, ** p<1×10^-4^, *** p<1×10^-7^. **(C)** smFISH of *luciferase* RNA in G3BP-GFP-λN transfected cells during arsenite stress. Scale bars = 2 µm. **(D)** smFISH of *luciferase* 25 BoxB RNA in G3BP-GFP-λN transfected cells during arsenite stress. Scale bars = 2 µm.

A key observation was that upon mild to moderate G3BP1-GFP-λN expression, *luciferase* RNAs with 7 or 25 λN binding sites shifted *luciferase* from a median of 14% partitioning into SGs to ∼36% or ∼62% partitioning, respectively (**Figure 2B-D**). Notably, this effect is dose-dependent since mRNAs with 25 BoxB sequences showed increased SG partitioning as compared to mRNAs with 7 BoxB sequences (**Figure 2B**). Addition of BoxB repeats to *luciferase* did not affect RNA partitioning to SGs when expressed without any λN protein or when co-expressed with GFP-λN (**Figure 2B**). Thus, the observed effect is due to the ability of the transcript to interact with G3BP1 and is not simply due to the increased length of the transcript. Examination of individual transfected cells showed that the reporter partitioning into SG was similar over a range of G3BP1 concentration (**Figure S3**), suggesting the effect is not a result of G3BP1 overexpression. Cells that highly over-expressed G3BP1, which leads to the formation of constitutive SG (Tourrière et al., 2003), were not included in this analysis. Taken together, these observations demonstrate that, at least during arsenite stress, the presence of multiple G3BP1 proteins on an mRNP can alter the partitioning of that mRNA to SGs in a dose-dependent manner. Therefore, at least some SG RBPs can influence the partitioning of client RNAs to SGs.

To determine if this effect was unique to G3BP1-GFP-λN, we also tethered GFP-λN or Halo-λN tagged versions of the stress granule components TIA1 or FMRP, and a G3BP1-Halo control to *luciferase* mRNAs with 25 BoxB sites. We observed that TIA1, but not FMRP, could recruit the reporter mRNA into SGs to the same extent as G3BP1 (**Figure 3**). This observation suggests that some SG-enriched RBPs can actively recruit client RNAs to SGs while other RBPs may simply enrich in SGs due to their association with SG-enriched transcripts.

**Figure 3:**
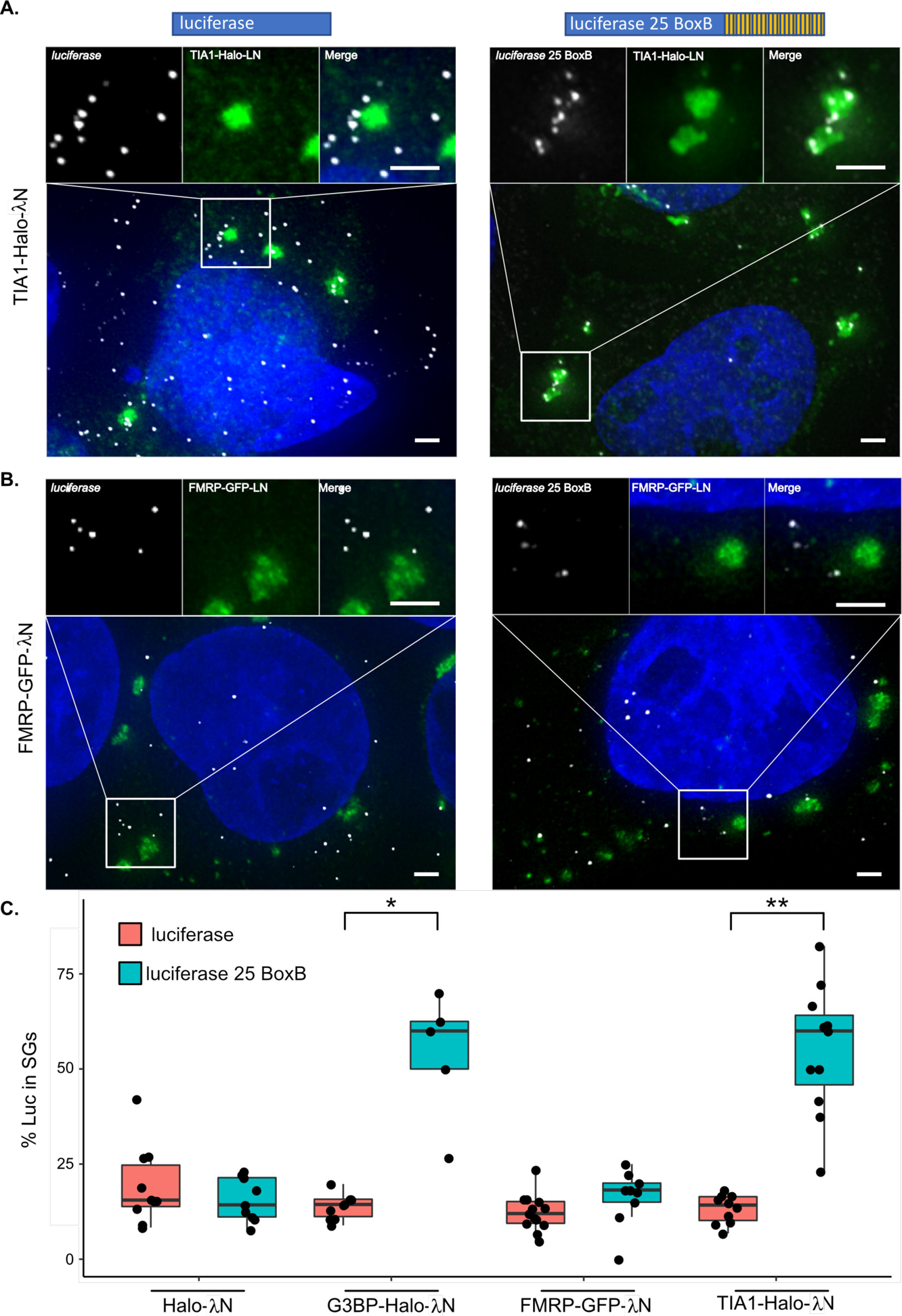
Tethering of TIA1, but not FMRP, increases *luciferase* RNA enrichment in SGs (A) smFISH of *luciferase* and *luciferase* 25 BoxB RNA co-expressed with TIA1-Halo-λN. Scale bars = 2 µm. **(B)** smFISH of *luciferase* and *luciferase* 25BoxB RNA co-expressed with FMRP-GFP-λN. Scale bars = 2 µm. **(C)** Boxplot depicting enrichment of *luciferase* 0 BoxB, and 25 BoxB RNA when tethered to G3BP, FMRP, and TIA1 through λN-BoxB interactions. *p<0.01, **p<1×10^-5^

### Quantitative estimation of the number of SG interactions with mRNAs

Based on our G3BP1 tethering experiments, we performed mathematical modeling of SG recruitment as a function of the number of RNA-RBP interactions. In our modeling, we assume all BoxB sites are occupied by G3BP-GFP-λN. Thus, the starting *luciferase* reporter mRNA has an unknown number of interactions, *n*, with SGs, yielding 14% SG partitioning. The mRNA *luciferase* reporter with 7 BoxB sites has *n*+7 interactions, yielding 36% SG partitioning, and the *luciferase* reporter with 25 BoxB sites has *n*+25 interactions, yielding 62% SG partitioning. We also assume an RNP with no interactions with SG would show limited enrichment in SGs (1% SG partitioning).

We performed curve fitting analysis allowing *n* to vary from 1 to 8 *luciferase* interactions with SG RBPs prior to the addition of any BoxB sequences. Curve fitting yielded a logarithmic fit (y = 18.16ln(x) - 0.08) that showed a maximum R^2^ value of ∼0.99 for the fit in which n = 2 (**Figure 4A, B**). It should be noted that n = 2 is only an estimate of the number of “G3BP-SG equivalent interactions”, which we define by the contribution of a tethered G3BP1 to the partitioning of *luciferase* mRNA into SGs and does not necessarily correspond to two discrete molecular interactions. However, similar conclusions are reached if the analysis below is repeated with any value of n between 1 and 8. Importantly, the shape of the fit raised several important implications.

**Figure 4:**
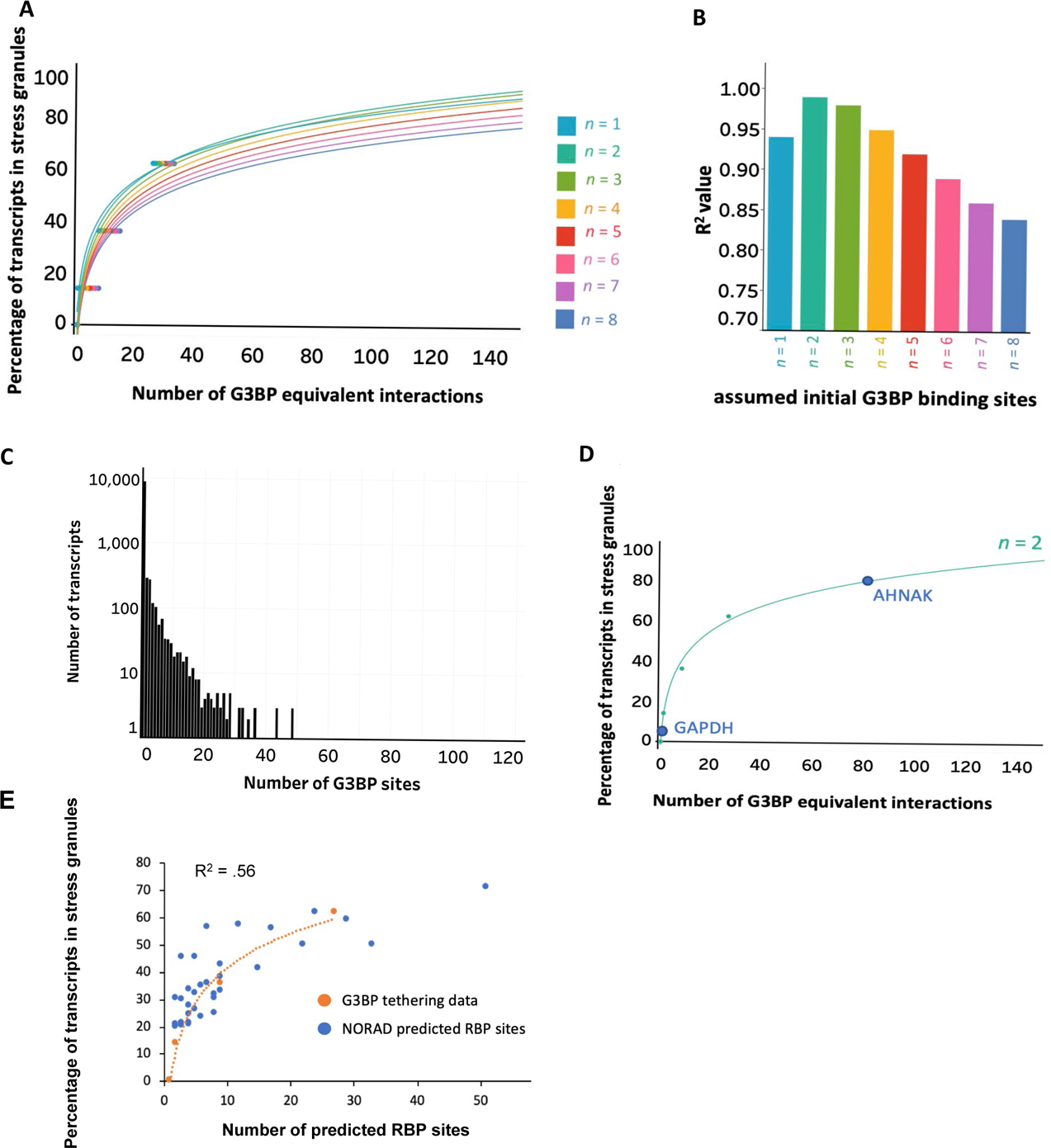
Mathematical modeling of RBP interactions suggest deletion of G3BP may have a limited effect on RNA enrichment in SGs **(A)** Curve fitting of the percentage of *luciferase* transcripts in SGs vs. number of G3BP-SG equivalent interactions. The predicted number of starting SG-interactions *luciferase* has before addition of BoxB sequences is represented by n. **(B)** Bar graph showing how the R^2^ value from curve fitting analysis changes with respect to n. **(C)** Histogram depicting the number of transcripts that contain various numbers of G3BP eCLIP peaks. Note that y-axis is on a log scale. **(D)** Prediction of the number of average “G3BP-SG equivalent interactions” of GAPDH and AHNAK based on their percent localization to SGs. **(E)** Curve fitting overlaying the predicted summation PRE, ARE, and SAM68 sites for NORAD fragment transcripts and G3BP tethering experiments.

First, this result suggests that the same RNP interactions with a SG will not affect the recruitment of all transcripts to SGs equally. For example, loss of a single SG interaction from an mRNP with 27 total “G3BP-SG equivalent interactions” would only reduce the expected proportion of mRNAs in SGs from 60% to 59%. In contrast, the loss of one interaction for an mRNP with three “G3BP-SG equivalent interactions” should reduce its accumulation in SGs from ∼20% to ∼13%. These observations highlight the key point that RNPs with small numbers of interactions with SGs should be affected to a greater extent by changing a single interaction, while RNPs with large numbers of SG interactions should be less affected by loss of a single RNA-RBP interaction.

Second, the maximum effect of a given RBP-RNA interaction on SG partitioning should be relatively small, and much less for RNAs which have many non-redundant RNA-RBP interactions. This second observation is important because eCLIP analysis shows that the majority of transcripts in the cell have 0-3 G3BP binding sites (Van Nostrand et al. 2018, **Figure 4C**).

Third, assuming all SG-targeting interactions are similar, one can make an estimate for the number of interactions a given mRNA has with SGs (**Figure 4D**). We used the *n*=2 curve, since it yielded the curve with the best fit (**Figure 4B**). From this curve, we made a transcriptome-wide first approximation of the number of interactions a given mRNA forms with SGs (**Supplemental Table S2**). For example, we would predict that *AHNAK*, where a median of 81% of the molecules are in SGs, has ∼87 “G3BP-SG equivalent interactions” with SGs (**Figure 4D**). Conversely, *GAPDH*, with a median of 3.4% of its molecules in SGs, is predicted to have ∼1 “G3BP-SG equivalent interactions” with SGs (**Figure 4D**). Given the assumptions in this model, these numbers should only be taken as a first approximation of interactions that will need to be refined in future analyses.

One assumption in this modeling is that every BoxB site is directly interacting with a G3BP1 molecule. If we assume that only half of the sites are occupied at a given time, then our estimates would necessarily be an overestimate of the number of interactions a given transcript forms with SGs (**Figure S4**). Still, the logic remains valid that the higher the number of interactions an RNA can form with SGs, the less sensitive that RNA will be to perturbations of individual interactions.

### Modeling Explains Behavior of the NORAD-luciferase constructs

To see if we can understand the *NORAD-luciferase* chimeric RNA results in terms of different numbers of RBPs, we first estimated the number of proteins bound to each region (**Supplemental Table S1**). In this analysis, we counted PREs, which serve as binding sites for Pumilio (Lee et al., 2016; Tichon et al., 2016), AREs, which can bind a diverse number of different ARE binding proteins and have been shown to correlate with SG enrichment (Namkoong et al., 2018), and Sam68 binding sites, which have also been mapped to NORAD (Tichon et al., 2018). We then plotted the number of predicted RBPs bound to each construct versus the SG enrichment. Remarkably, this set of data points fit the curve derived from the tethered G3BP1 RNAs quite well (**Figure 4E**). Moreover, this analysis would predict that the PRE mutant, which has 18 deleted RBP binding sites, should change the recruitment of the *luciferase* reporter from 71% to 61%, which is similar to the observed 50%. The deviation between recruitment of the PRE mutant and the predicted value may be due to the PRE mutations disturbing other protein binding sites. Thus, the behavior of both the tethered G3BP1 reporters, and the *luciferase-NORAD* chimeric mRNAs, can be explained as being due to the summation of multiple interactions that together increase the partitioning of RNPs into SGs.

We also estimated the number of RBPs bound to *NORAD* segments based on CLIP sites from a database containing a meta-analysis of RBP CLIP studies (Yang et al.,2015, **Figure S5A**). We found that SG RBP CLIP sites showed a qualitative correlation with the enrichment of individual *NORAD* segments within SGs (**Figure S5B**), and that when we summed the total number of SG RBPs for each segment of *NORAD* there was a good correlation between SG CLIP sites and *NORAD* segment enrichment (**Figure S5C**). Taken together, these results are consistent with a model in which RBPs can act in tandem to define the RNA composition of SGs.

### Loss of G3BP1&2 does not globally affect RNA localization to SGs

Our modeling of SG-mRNA interactions suggests that RBPs act in tandem to contribute to RNP enrichment within SGs. If this model is true, we would anticipate that deletion of individual SG RBPs, such as G3BP, should have a limited effect on the recruitment of transcripts. To test this prediction, we desired to purify and determine the SG transcriptome from cells with and without G3BP1/G3BP2. In order to perform this experiment, we needed to purify SGs using a different SG component than G3BP1, which was used in earlier work (Khong et al., 2017), and under sorbitol stress condition, where G3BP1 and G3BP2 are not required for SG formation (Kedersha et al., 2016). Thus, we first tested whether immunopurification using an antibody to the SG component PABPC1 yielded a similar SG transcriptome as that seen with GFP-G3BP1 under arsenite stress (Khong et al., 2017; **Figure 5A-B**).

**Figure 5:**
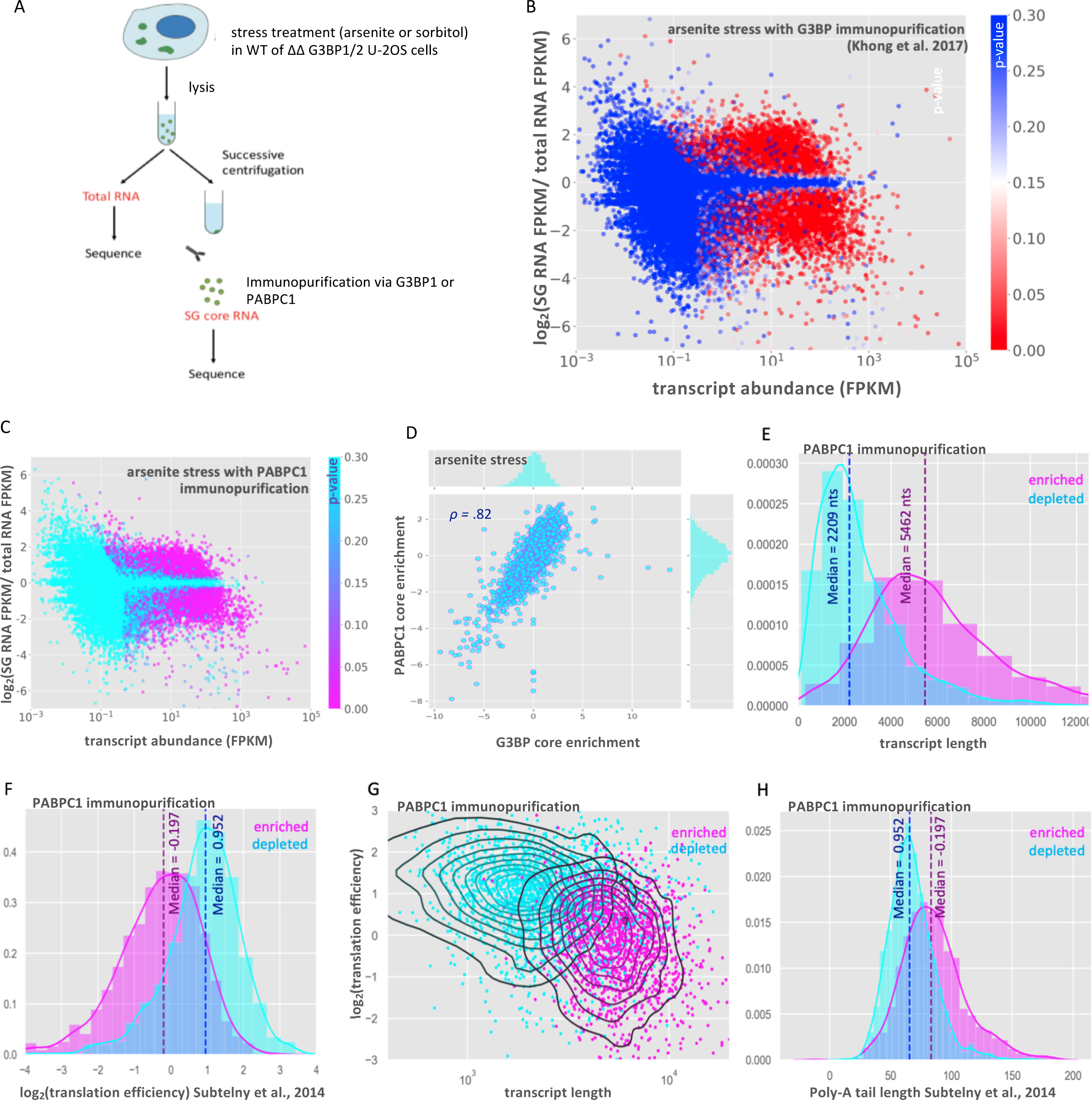
RNA-Seq of the PABPC1 stress granule core transcriptome **(A)** Schematic of SG core purification strategy in WT or ΔΔG3BP1/2 cells treated with either arsenite or sorbitol stress and using G3BP1 or PABPC1 for immunopurificaiton. **(B)** MA-plot showing stress granule enrichment vs. abundance from previous work examining the SG transcriptome via G3BP1 immunopurification during arsenite stress (Khong et al., 2017). **(C)** MA-plot showing stress granule enrichment vs. abundance from PABPC1 immunopurification during arsenite stress. **(D)** Scatterplot of RNA enrichment in PABPC1 cores vs. enrichment in G3BP cores during arsenite stress. **(E)** Histogram showing distribution of SG-enriched and SG-depleted transcripts with respect to transcript length (purified with PABPC1). **(F)** Same as E but for translation efficiency. **(G)** Scatterplot of translation efficiency vs. transcript length with kernel density estimate overlay. Color-coded by enrichment/depletion in SGs. **(H)** Same as E, but for poly-A tail length.

Total RNA and SG core purification via PABPC1 pulldown yielded reproducible transcriptomes (**Figure S6**). SG RNA transcriptomes based on the purification of SG cores with PABPC1 antibody showed little similarity to total RNA transcriptomes, with 3251 transcripts enriched in SGs (p < 0.01), and 3693 transcripts depleted from SGs (p < 0.01) (**Figure 5C**). Comparison of the RNAs enriched in SG cores showed that enrichment scores from PABPC1-purified cores and GFP-G3BP1-purified cores had a strong linear correlation (**Figure 5D**, Pearson’s r = 0.82). Thus, PABPC1 pulldown identifies a population of RNAs that strongly overlaps with the SG transcriptome identified by GFP-G3BP1 pulldown. Consistent with PABPC1 and GFP-G3BP1 purification identifying similar RNAs, we observed that in both cases, RNAs enriched in SGs are biased towards long, poorly translated RNAs (Khong et al., 2017; Subtelny et al., 2014; **Figure 5E-G**). Additionally, we observe that enriched RNAs tend to have longer poly-A tails (data obtained from Subtelny et al., 2014), which could be explained by transcripts with longer poly-A tails having less efficient translation rates, lower overall abundance, or by the increased length of these transcripts, which would add additional RBP interaction elements (Lima et al., 2017, **Figures 5H, S7**). Taken together our findings suggest that PABPC1 and G3BP1 SG cores share a similar RNA composition and that the RNAs that co-purify with PABPC1 and G3BP1 cores have similar physical properties.

Since ΔΔG3BP1/2 cell lines form SGs during sorbitol treatment, we planned on analyzing SG transcriptomes from sorbitol-treated cells with and without G3BP1/2 expression. Before sequencing PABP-containing SGs from WT and ΔΔG3BP1/2 cells during sorbitol stress, we first examined whether the RNA composition of sorbitol-induced SGs was similar to arsenite-induced SGs. Thus, we compared the transcriptome of SG cores purified via PABPC1 and GFP-G3BP1 immunopurification during hyperosmotic stress induced by sorbitol (Dewey et al., 2011; Kedersha et al., 2016; **Figures 6, S8, S9**).

**Figure 6:**
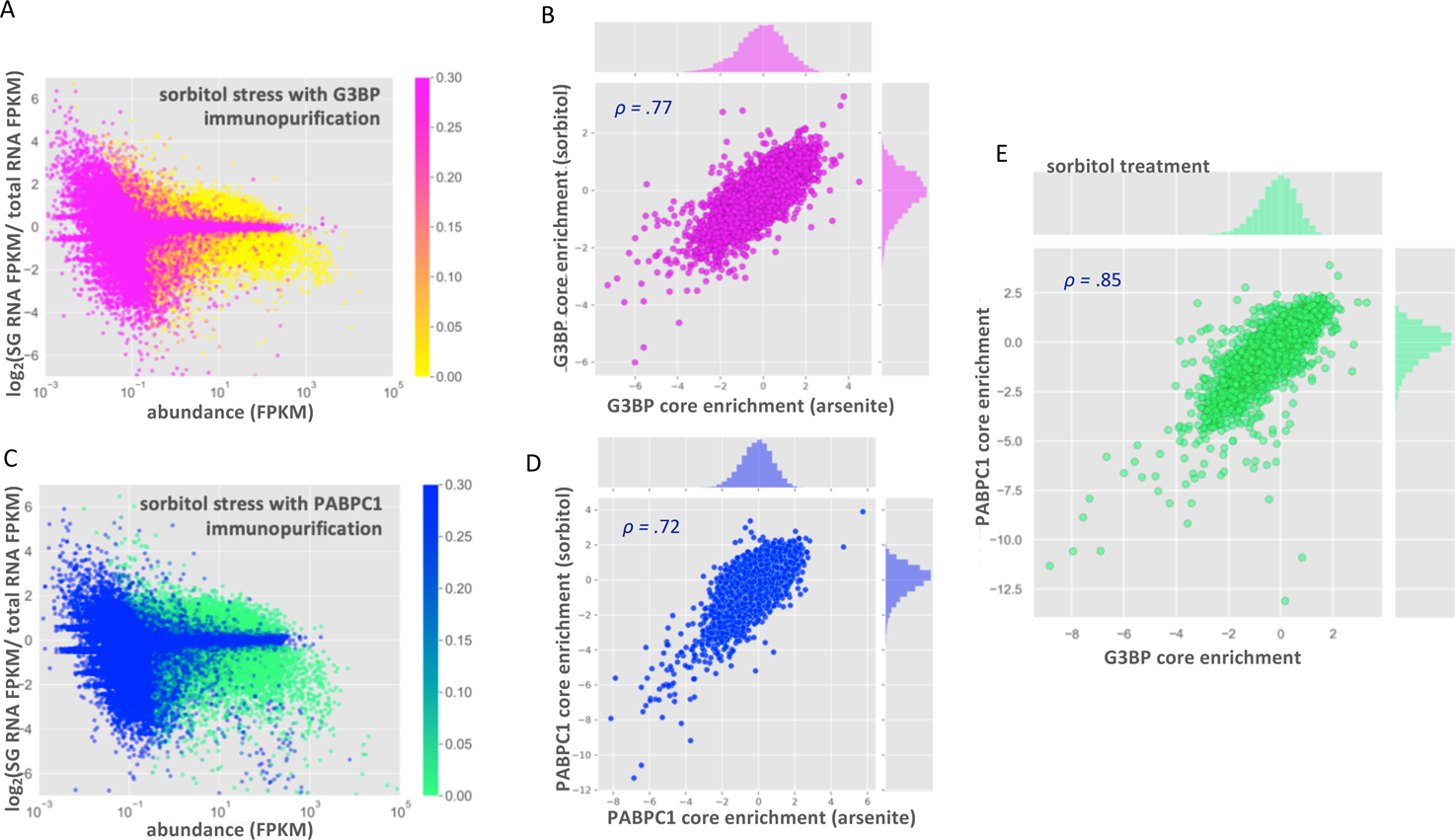
RNA localization to stress granule cores is conserved between sorbitol and arsenite stress **(A)** MA-plot showing stress granule enrichment vs. abundance for G3BP1 purified SGs during sorbitol treatment. **(B)** Scatterplot showing correlation between G3BP core enrichment scores in sorbitol vs. arsenite stress. **(C)** MA-plot showing stress granule enrichment vs. abundance for PABPC1 purified SGs during sorbitol treatment. **(D)** Scatterplot showing correlation between RNA enrichment scores from PABPC1 core purification in sorbitol vs. arsenite stress. **(E)** Scatterplot showing correlation between enrichment scores from PABPC1- and G3BP-purified SG cores during sorbitol stress.

Purification of GFP-G3BP1 SGs under sorbitol stress yielded a very similar transcriptome to that observed under arsenite stress, with 2829 significantly enriched and 3721 significantly depleted transcripts (**Figure 6A-B**). This is in agreement with the previous observations that the SG transcriptome is conserved between many different stresses (Khong et al., 2017; Namkoong et al., 2018). We also observed that the SG transcriptome based on PABPC1 immunopurification is largely conserved between arsenite and sorbitol stresses and is similar to the SG transcriptomes defined by G3BP1 immunopurification (**Figure 6C-E**). Taken together, our results indicate that the SG transcriptome is highly similar between arsenite and sorbitol stress conditions, and that the mRNAs pulled down are largely independent of the SG protein used for the affinity purification.

We then examined whether G3BP1 and G3BP2 affected the mRNAs partitioning into SGs during sorbitol stress. To test this possibility, we purified sorbitol-induced SG cores from WT and ΔΔG3BP1/2 U-2 OS cells using antibodies to PABPC1 (**Figures S9, S10**). Strikingly, sorbitol-induced SGs contained a similar transcriptome regardless of whether cells expressed or lacked G3BP1 & G3BP2 (**Figure 7A, B**). The enrichment scores for these transcriptomes showed a strong linear correlation (R = 0.94), suggesting that the enrichment of RNA in SGs is largely independent of G3BP during sorbitol stress. We interpret this observation to argue that G3BP1 and G3BP2 do not generally affect the mRNAs recruited to SGs.

**Figure 7:**
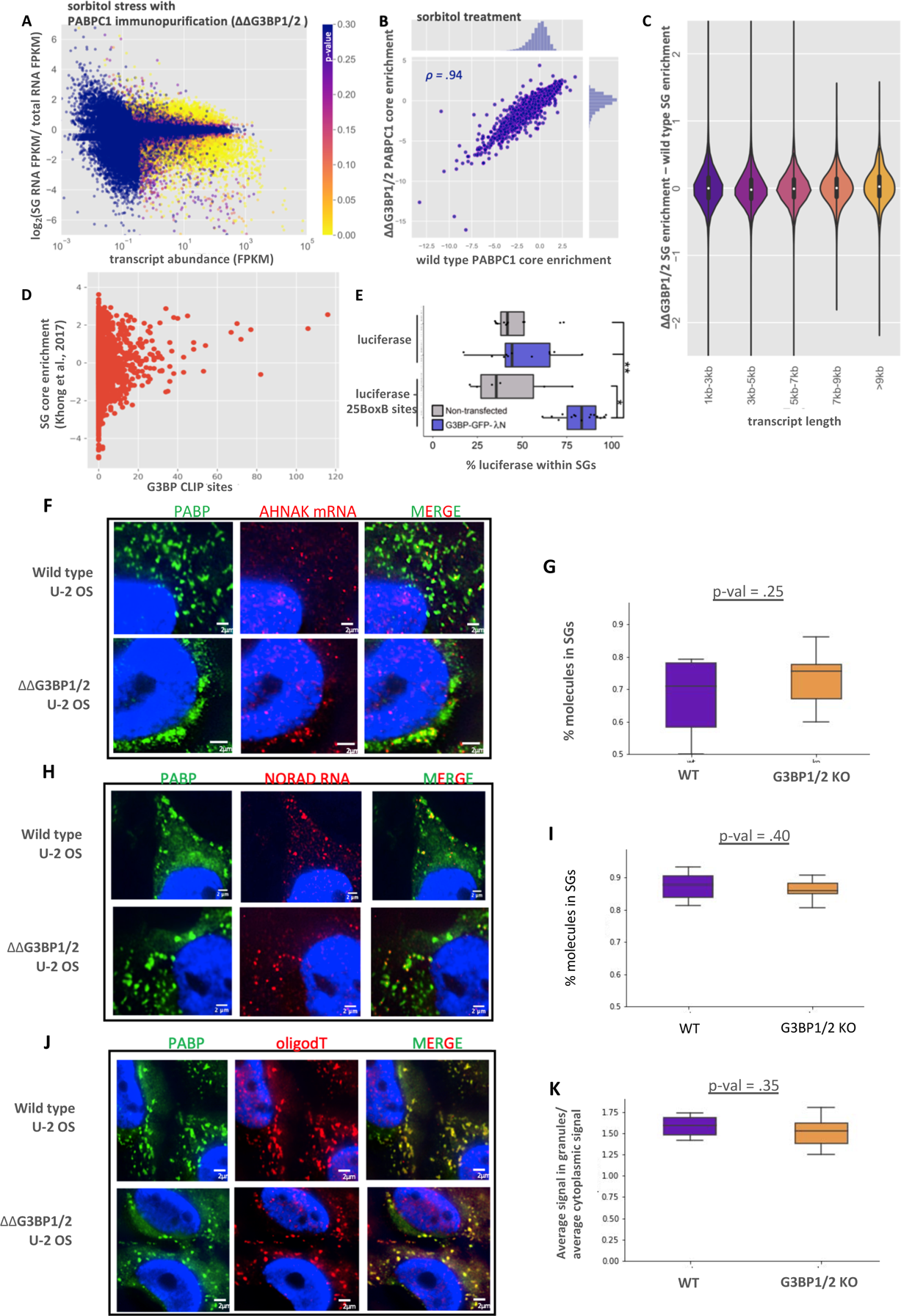
Global RNA localization to stress granule cores is independent of G3BP **(A)** MA-plot showing stress granule enrichment vs. abundance for PABPC1 purified SGs during sorbitol treatment in ΔΔG3BP1/2 cells. **(B)** Scatterplot depicting the enrichment of transcripts in PABPC1 stress granule cores in ΔΔG3BP1/2 vs WT U-2 OS cells during sorbitol stress. **(C)** Violin plots depicting ΔΔG3BP1/2 - WT stress granule enrichment scores for all transcripts, binned by length. **(D)** Scatterplot depicting the fraction of a transcript that is localized to SGs during arsenite stress vs. the number of G3BP eCLIP peaks for that transcript. **(E)** Boxplot showing the localization to SGs of *luciferase* reporter RNAs with 0 or 25 BoxB sites during sorbitol stress in non-transfected and G3BP-GFP-λN transfected cells. * p<0.01, ** p<0.001. **(F)** smFISH of *AHNAK* mRNA during sorbitol stress in WT and ΔΔG3BP1/2 cells. **(G)** Boxplot quantifying the fraction of *AHNAK* cytoplasmic transcripts in SGs in WT and ΔΔG3BP1/2 cells**. (H)** smFISH of *NORAD* mRNA during sorbitol stress in WT and ΔΔG3BP1/2 cells. **(I)** Boxplot quantifying the fraction of *NORAD* in SGs in WT and ΔΔG3BP1/2 cells**. (J)** FISH of poly(A) RNA during sorbitol stress in WT and ΔΔG3BP1/2 cells. **(K)** Boxplot quantifying average intensity of poly(A) signal inside SGs vs. average intensity of poly(A) signal in the cytoplasm.

Interactions between G3BP and mRNA might act in concert with other RBP-RNA, or RNA-RNA interactions to drive RNAs into SGs (Van Treeck et al., 2018; Khong et al., 2017). In this view, G3BP would only affect the recruitment of mRNAs to SGs that were on the cusp of SG enrichment. In order to examine this possibility, we calculated the change in ΔΔG3BP1/2 SG enrichment scores vs. WT enrichment scores as a function of RNA length. If G3BP acted in concert with other RNA-RNA and RNA-RBP interactions to drive localization to SGs, one would anticipate that G3BP would only have an observable effect for transcripts of shorter lengths, which have fewer interactions. However, we observed similar SG enrichment between WT and ΔΔG3BP1/2 cell lines for RNAs of any length (**Figure 7C**). Thus, we observed no significant impact of G3BP1 and G3BP2 on mRNA targeting to SGs even when binning for different lengths of mRNAs.

We reasoned that only transcripts that exhibit G3BP binding capability would show differential recruitment to SGs. Thus, we utilized data from a recent eCLIP study which examined G3BP eCLIP targets (Van Nostrand et al., 2016). However, we observed that G3BP binding as assessed by eCLIP and SG enrichment were only weakly positively correlated (**Figure 7D**). Moreover, G3BP target transcripts showed no altered localization in ΔΔG3BP1/2 cells, nor when we limited the analysis to short transcripts (**Figure S11**).

Since G3BP is required for SG formation during arsenite stress, but not sorbitol stress, it is possible that G3BP simply does not influence the partitioning of RNAs during sorbitol stress. To test this possibility, we examined if tethered G3BP1 could target the *luciferase* mRNA to SGs under sorbitol stress in a manner similar to what was observed under arsenite stress. We observed that G3BP1 tethering led to a significant and reproducible increase in *luciferase* localization to SGs during sorbitol stress (**Figure 7E**). Thus, G3BP1 can still artificially drive SG enrichment even in sorbitol stress wherein G3BP is not required for SG formation.

A final possibility for why the SG transcriptome does not change in the ΔΔG3BP1/2 cell lines is that depletion of G3BP affects all transcripts equally, which might be missed in our sequencing data. Thus, we examined whether global RNA recruitment to SGs was altered in ΔΔG3BP1/2 cells. By smFISH, we detected no discernable difference in the localization of *AHNAK* (**Figure 7F, G**). This is in spite of the fact that *AHNAK* has over 70 predicted G3BP CLIP sites (**Figure 7D**). This suggests that even transcripts with many G3BP sites may see modest changes in partitioning, likely due to compensating interactions that extremely large mRNAs like *AHNAK* (18 kb) are capable of forming. Both *NORAD* and total poly(A)+ signal showed a slight reduction in SG partitioning in ΔΔG3BP1/2 cells (**Figure 7H-K**), although these reductions did not achieve statistical significance. These findings argue that G3BP does not strongly affect the SG transcriptome. More broadly, this suggests that individual interactions between an mRNP and a SG that are of a similar strength as G3BP interactions with SGs will have minimal effects on mRNP partitioning into SGs. Therefore, RNA localization to SGs largely arises through the synergistic effects of multiple interactions acting in concert.

## DISCUSSION

In this work we present evidence that specific RNA-binding proteins can target mRNPs into SGs. The critical observation is that tethered G3BP1 or TIA1 can increase the partitioning of the *luciferase* mRNA into SGs (**Figures 2, 3**). Although it has been widely anticipated that RBPs can target mRNAs to SGs, to our knowledge this observation provides the first demonstration of this principle. Possible mechanisms by which mRNAs could be targeted to SGs by G3BP1 or TIA1 include the formation of G3BP1 dimers (Kedersha et al., 2016), or interactions between the TIA1 prion-like domain (Gilks et al., 2004).

We present two lines of evidence that the partitioning of an mRNP into SGs will be based on multiple elements acting in an additive manner. First, we observed that multiple elements within the *NORAD* RNA were sufficient to increase the SG enrichment of a *luciferase* reporter RNA (**Figure 1**). Second, we demonstrated that the ability of tethered G3BP1 to target the reporter mRNA to SGs was dose-dependent (**Figure 2**). The ability of protein interactions to target mRNPs to SGs suggests that the partitioning of an mRNP into a stress granule will be a summation of protein-protein, protein-RNA and RNA-RNA interactions between an individual mRNP and the SG. This is directly analogous to the hypothesis that SGs form through various combinations of interactions between different mRNPs (Van Treeck et al., 2018). Thus, the recruitment of mRNPs into SGs reflects the summation of a number of interactions between individual mRNPs.

The recruitment of RNPs can be considered a simple equilibrium binding reaction with the partitioning of the RNP being proportional to e^-ΔG/RT^. Moreover, we observed that the SG enrichment of individual RNPs was related to the number of interactions in a ln dependent manner (**Figure 4**). This is consistent with the model where the energetics of each individual interaction sum together to give an overall ΔG for SG partitioning as predicted by simple equilibrium binding.

By using some reasonable assumptions, we are able to make estimations of the number of interactions a given RNP will have with SGs and its partition coefficient. In this analysis, we estimate the number of interactions based on their similarity to a G3BP1 interaction. While this is undoubtedly an over-simplification, it allows for the first estimation of how many interactions RNPs have within SGs. Moreover, we note that tethered G3BP1 and TIA1 gave similar degrees of SG partitioning of the *luciferase* mRNA with 25 BoxB sites, suggesting that at least these two proteins have similar interactions with SGs. These estimates suggest that mRNAs with less than 20% of their mRNAs in SGs have 1-5 “G3BP-SG equivalent interactions”, while mRNAs that partition greater than 50% of their mRNAs in SGs have over 15 “G3BP-SG equivalent interactions”. Although these are crude estimates these numbers make two important points. First, the number of interactions a highly enriched RNP has in SGs is quite high. For example, we estimate that *AHNAK* mRNA will have 97 “G3BP-SG equivalent interactions”. This highlights the second important conclusion: any given interaction only makes a small contribution to the overall enrichment of mRNAs in SGs. For example, for the *AHNAK* mRNA we estimate removing a single interaction between the mRNP and SG would alter the enrichment by 0.2%.

As a test of this rationale, we examined the global RNA composition of SGs in cells lacking the highly abundant SG RBPs G3BP1 and G3BP2. In agreement with our hypothesis, we found that WT and ΔΔG3BP1/2 cell lines contain a similar SG transcriptome (**Figure 7**). This observation was confirmed by smFISH for *AHNAK*, *NORAD*, and by oligo(dT) staining.

Taken together, we propose a model in which multiple RBP-RNA and RNA-RNA interactions act synergistically to define the RNA composition of SGs (**Figure S12**). In this model, deletion or depletion of any single RBP would have a negligible effect on RNA localization to SGs. which is supported by the evidence that G3BP1 & G3BP2 deletions had essentially no effect on the SG transcriptome (**Figure 7**). Similarly, cells lacking Pumilio proteins can still accumulate *NORAD* in SGs, based on the sequencing of a heavy RNP containing fraction (Namkoong et al. 2018). This is likely due to the ability of other RBPs and RNA-RNA interactions to compensate for the absence of G3BP or Pumilio. The fact that the RNA composition of SGs is largely conserved between different stresses (Khong et al., 2017, Namkoong et al., 2018, **Figure 6**) also supports our estimations. Since some proteins are only recruited to SGs under specific stresses (Buchan and Parker, 2009; Markmiller et al., 2018), the loss of these potential interactions in SGs has little effect on the global RNA composition of SGs. Thus, there are likely multiple synergistic/redundant interactions that lead to RNA enrichment in SGs. This is not to say that individual RBPs cannot modulate the ability of an RNA to enrich in SGs. Indeed, increasing the number of RBPs on a transcript can lead to the enhanced enrichment of that transcript within SGs (**Figure 2, 3**). We hypothesize that other SG RBPs should also be able to modulate the recruitment of transcripts similarly to G3BP tethering. It remains possible that there will be some molecular interactions between RNPs and SGs that are much stronger than the average “G3BP-SG equivalent interaction” we define here and will therefore have a larger effect on the SG partitioning of an individual mRNP.

An apparent conundrum from these observations is that G3BP1 is required for stress granule formation in many stresses, but its absence does not notably alter the transcriptome of SG. We suggest that the resolution to this conundrum is due to the fundamental differences in the kinetics of stress granule assembly as compared to the recruitment of individual mRNPs into SGs. The formation of SGs is a cooperative process wherein the average interaction between mRNPs will set a critical threshold above which mRNP nucleation into SGs will initiate, followed by recruitment of additional mRNPs to form SG cores. If the average interaction between mRNPs is altered even by 10% this can shift the system into a region of non-assembly due to the highly cooperative nature of SG cores, which contain 20-70 mRNAs (Khong et al., 2017). Thus, SG assembly is highly cooperative and very sensitive to the average interaction strength between individual mRNPs. Such highly cooperative assembly, and sensitivity to small changes in average interactions is a general property of any large assembly made up of multiple components, which provides numerous opportunities for the regulation of higher scale assembly. In contrast, the recruitment of an individual mRNP into an existing SG can be considered a simple equilibrium binding reaction without a cooperative effect of changes in mRNP affinities such that changes in K_D_s have modest effects on mRNA partitioning.

This analysis makes several predictions. First, components required for SG formation can halt overall assembly without substantially affecting the transcriptome of SGs in cases where they do form. Second, increasing SG formation artificially (e.g. by inhibiting eIF4A (Tauber et al., 2020)); or by overexpressing TIA1 will still generally lead to the same mRNAs accumulating in SG. Third, genetic or pharmacological manipulations that alter the initiation events, or average interactions between mRNPs, even by a small amount, can prevent SG formation.

## METHODS

### Plasmid Construction

For a list of plasmids constructed for use in this manuscript, as well as oligos used for plasmid construction, please see **Supplemental Table S3**. Tet-inducible Luciferase reporter was a gift from Moritoshi Sato (Addgene plasmid # 64127; http://n2t.net/addgene:64127; RRID:Addgene_64127, Nihongaki et al., 2015). BoxB repeats were cloned out of pCMV5-25BoxB, which was a gift from Maria Carmo-Fonseca (Addgene plasmid # 60817; http://n2t.net/addgene:60817; RRID:Addgene_60817, Martin et al., 2013) and placed into the 3’UTR of the luciferase reporter. Vector used as the backbone for AAVS targeting of Cas9 was pRP2855, an AAVS-TDP43 plasmid, in which the TDP43 was excised by restriction digest and replaced with *luciferase* constructs. G3BP-GFP-λN and GFP-λN plasmids were gifts from Richard Lloyd’s lab. FMRP was cloned out of pFRT-TODestFLAGHAhFMRPiso1, a gift from Thomas Tuschl (Addgene plasmid # 48690; http://n2t.net/addgene:48690: RRID:Addgene_48690, Ascano et al., 2012). NORAD-PREmut plasmid was a gift from Josh Mendel’s lab and placed into the 3’ UTR of the *luciferase* reporter.

## Genomic integration into AAVS locus

Cells were transfected with 1 μg CRISPR/Cas9 plasmid (pRP2854) in conjunction with 1 μg appropriate *luciferase* reporter construct (pRP2856, pRP2873 and pRP2874) using Jet Prime reagent. Transfection of pRP2854 alone was used as a negative control. 24 hours following transfection, cells were split from a 6-well plate to a 10 cm dish. After another 24 hours, media was replaced with new media containing 1 μg/mL puromycin to begin selection for cells with genomic integration. Following 24 hours with puromycin selection, media was replaced with fresh puromycin. After all cells were dead in the negative control plate, media was replaced with fresh media lacking puromycin for 48 hours. This is an optional step to help get rid of any residual plasmid. Puromycin was then added for another 48 hours to finalize the selection. Single colony selection was not done for these experiments.

## Stellaris smFISH probes

Custom Stellaris smFISH probes against *AHNAK*, *NORAD* and firefly *luciferase* transcripts were designed with Stellaris RNA FISH Probe Designer (Biosearch Technologies, Petaluma, CA), available online at http://www.biosearchtech.com/stellaris-designer (version 4.2). AHNAK and *NORAD* smFISH probes, labeled with Quasar 670 dye, and firefly *luciferase* probes, labeled with Quasar 570, were ordered from Stellaris (Biosearch Technologies, Petaluma, CA).

## Sequential IF and FISH

Sequential immunofluorescence and smFISH on fixed U-2 OS cells was performed with Stellaris buffers or homemade buffers (Dunagin et al., 2015) according to the manufacturer’s protocol: (https://biosearchassets.blob.core.windows.net/assets/bti_custom_stellaris_immunofluorescence_seq_protocol.pdf).

Briefly, U-2 OS cells were seeded on sterilized coverslips in 6-well tissue culture plates. At ∼80% confluency, media was exchanged 1 hour before experimentation with fresh media. After stressing cells (see stress conditions), the media was aspirated and the cells were washed with pre-warmed 1×PBS. The cells were fixed with 500 μL 4% paraformaldehyde for ten minutes at room temperature. After fixation, cells were washed twice with 1×PBS, permeabilized in 0.1% Triton X-100 in 1×PBS for five minutes and washed once with 1×PBS.

For IF detection, coverslips were incubated in primary antibody for 1 hour. Coverslips were washed three times with 1×PBS for 10 minutes each wash. Then cells were incubated in secondary antibody (Thermo Fisher Scientific A-31553). Again, coverslips were washed three times with 1×PBS for 10 minutes each wash. Then, cells were treated with smFISH Buffer A for 5 min. Coverslips were transferred to a humidifying chamber with smFISH probes and placed in the dark at 37°C for 16 hours. Coverslips were placed in Buffer A for 30 minutes in the dark, washed with Buffer B for 5 minutes and placed onto a slide with VECTASHIELD Antifade Mounting Medium with DAPI (Vector Labs, H-1200). For assays requiring quantification of smFISH probes in stress granules, VECTASHIELD Antifade Mounting Medium without DAPI was used (Vector Labs, H-1000).

In order to maintain consistency, the same protocol was utilized in IF only experiments, however the portions of the protocol calling for smFISH were omitted. Antibodies that were used include PABP (Abcam ab21060), G3BP (Abcam ab56574). In all imaging experiments at least 10 cells were imaged.

smFISH probes were labeled based on a recent protocol (Gaspar et al., 2017). Oligonucleotides used for smFISH were designed using the Stellaris Probe Designer (https://www.biosearchtech.com/support/education/stellaris-rna-fish). Briefly, oligos were labeled by incubating 4 μL of 200 μM pooled oligonucleotides (Integrated DNA Technologies), 1 μL of TdT (Thermo Fischer Scientific EP0161) and 6 μL of ddUTP (Axxora JBS-NU-1619-633) fluorophore for eight hours. After 8 hours, another 1 μL of TdT enzyme was added and the reaction was allowed to continue overnight. Probes were then ethanol precipitated by adding 164.5 μL of nuclease free water, 0.5 μL of 0.5 mg/ml linear acrylamide, 20 μL of 3 M sodium acetate (pH 5.5) and 800 μL of pre-chilled 100% ethanol. This mixture was then placed at −80°C for at least 20 minutes. Labeled oligos were pelleted by centrifugation at 16,000 ×g at 4°C. Probes were then washed 3 times by adding 1 mL of 80% ethanol and centrifugation at 16,000 ×g at 4°C. After washing, the pellet was allowed to air dry for 10 minutes and probes were resuspended in 25 μL of nuclease free water.

## Microscopy

Fixed U-2 OS cells stained by immunofluorescence and smFISH, were imaged using a wide field DeltaVision Elite microscope (Applied Biosystems) with a 100× objective and a PCO Edge sCMOS camera. At least five images with 20 Z-sections were taken for each experiment. All images in the manuscript are processed by FIJI (Schindelin et al., 2012) or Imaris (Bitplane).

## Imaris identification of smFISH spots

To measure the fraction of smFISH spots in stress granules, deconvolved images were analyzed using Bitplane Imaris image analysis software as described previously (Khong et al., 2018).

## U-2 OS growth conditions and reagents

Human osteosarcoma U-2 OS cells expressing G3BP1-GFP (Paul Taylor Lab), U-2 OS cells and U-2 OS ΔΔG3BP1/2 (Kedersha et al., 2016) were used in all experiments. All cells were maintained in DMEM with high glucose, 10% fetal bovine serum, and 1% penicillin/streptomycin at 37°C/ 5% CO_2_.

## Isolation of RNA from U-2 OS cells and SG cores for RNA-sequencing

Parental and ΔΔG3BP1/2 U-2 OS cells expressing G3BP1-GFP were grown to 85% confluency in three 500 cm^2^ TC-treated culture dishes (Thermo Fisher Scientific, 07-200-599). One hour prior to stress, cell culture media was exchanged with fresh media. Cells were then stressed with either NaAsO_2_ or sorbitol (see ‘Stress Conditions’). After stress, cells were washed once with media, transferred to falcon tubes, and pelleted at 1,500 ×g for 3 mins. Upon aspirating the media, the pellets were immediately flash-frozen in liquid N_2_ and stored at −80°C until isolation of mammalian SG cores was performed. PABPC1 SG cores were purified as previously described for the purification of G3BP cores, however instead of using anti-GFP for the pulldown of SG cores, 20 μL of anti-PABPC1 (ab21060) was used (Khong et al., 2017, 2018). For all experiments, biological triplicates were acquired for both SG-purified RNA and total RNA except in the case of PABPC1 SG RNA purified during sorbitol stress, in which duplicates were acquired (One of the replicates was excluded because it showed little similarity to the other biological replicates).

## Stress conditions

To examine mRNA localization during stress, we used the following stress conditions. For arsenite stress experiments, cells were treated with 0.5 mM sodium arsenite (Sigma-Aldrich S7400) for 1 hour. For osmotic stress, cells were stressed in 0.5M D-sorbitol (Sigma-Aldrich S1876) for 2.5 hours. Cells were fixed after the completion of each stress with 4% paraformaldehyde (Fisher Scientific NC0179595).

## Library construction and RNA-sequencing

RNA quality was assessed by TapeStation analysis at the Biofrontiers Institute Sequencing Facility. Paired-end cDNA libraries were prepared at the Biofrontiers Institute Sequencing Facility using the KAPA HyperPrep with RiboErase. cDNA libraries were sequenced on a NextSeq High output 150 cycle (2×75).

## Sequencing data analysis

Read quality was assessed using fastqc. An index genome was acquired from GENCODE (Release 19 GRCh37.p13). Reads were aligned using hisat2. Differential expression analysis was performed using Cuffdiff (version 2.2.1) with the default parameters (Trapnell et al., 2013). Gene Ontology analysis was performed using the gene ontology consortium (http://www.geneontology.org/). Transcript lengths were acquired from Ensembl’s Biomart Tool. All sequencing data can be found at NCBI GEO GSE119977.

## ACKNOWLEDGEMENTS

We would like to thank Dr. Amber Sorenson, Katelyn Hammond, and Caitlin Poling at the Biofrontiers Next Generation Sequencing Facility for the preparation and sequencing of RNA-Seq libraries. We would like to thank Matthew Deater and Richard Lloyd for the G3BP-GFP-λN and GFP-λN plasmids. Thanks to Josh Mendel for the NORAD-PRE plasmid. We would like to thank Nancy Kedersha and Paul Anderson for the ΔΔG3BP1/2 U-2 OS cell line as well as WT U-2 OS cells. The data analysis and visualization work was performed at the BioFrontiers Institute Advanced Light Microscopy Core. The Analysis Workstation and the software package Imaris were supported by NIH 1S10RR026680-01A1. We would also like to thank Paul Taylor for the G3BP1-GFP U-2 OS cells. This work was funded by the NIH (GM045443) and the Howard Hughes Medical Institute. B.V.T was supported by NSF SCR Training Grant T32GM08759.

## COMPETING INTERESTS

The authors declare that no competing interests exist.

---

**Figure S1:**
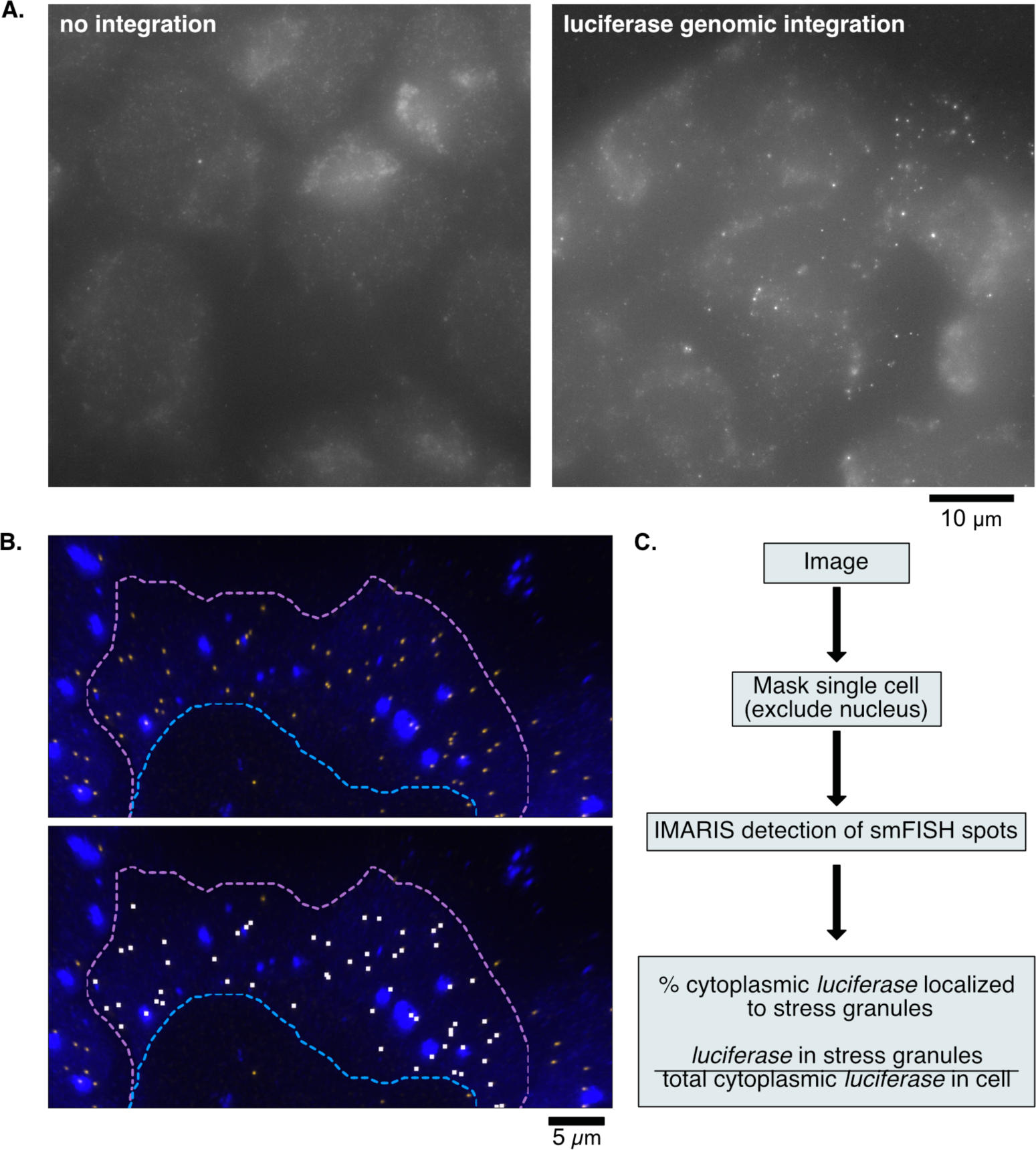
*Luciferase* as a reporter mRNA **(A)** *Luciferase* mRNA is not endogenous to mammalian cells. Non-deconvolved microscopy images showing smFISH signal in U-2 OS cells lacking genomic integration and cells with successful genomic integration. **(B)** *Luciferase* RNA is not enriched in SGs. Stress granules in blue, *luciferase* RNA in yellow (above) or demarcated by IMARIS software in white (below). **(C)** Workflow for smFISH quantification. Following imaging, single cells can be masked using IMARIS. From here, the percent of cytoplasmic *luciferase* localized to stress granules is calculated.

**Figure S2:**
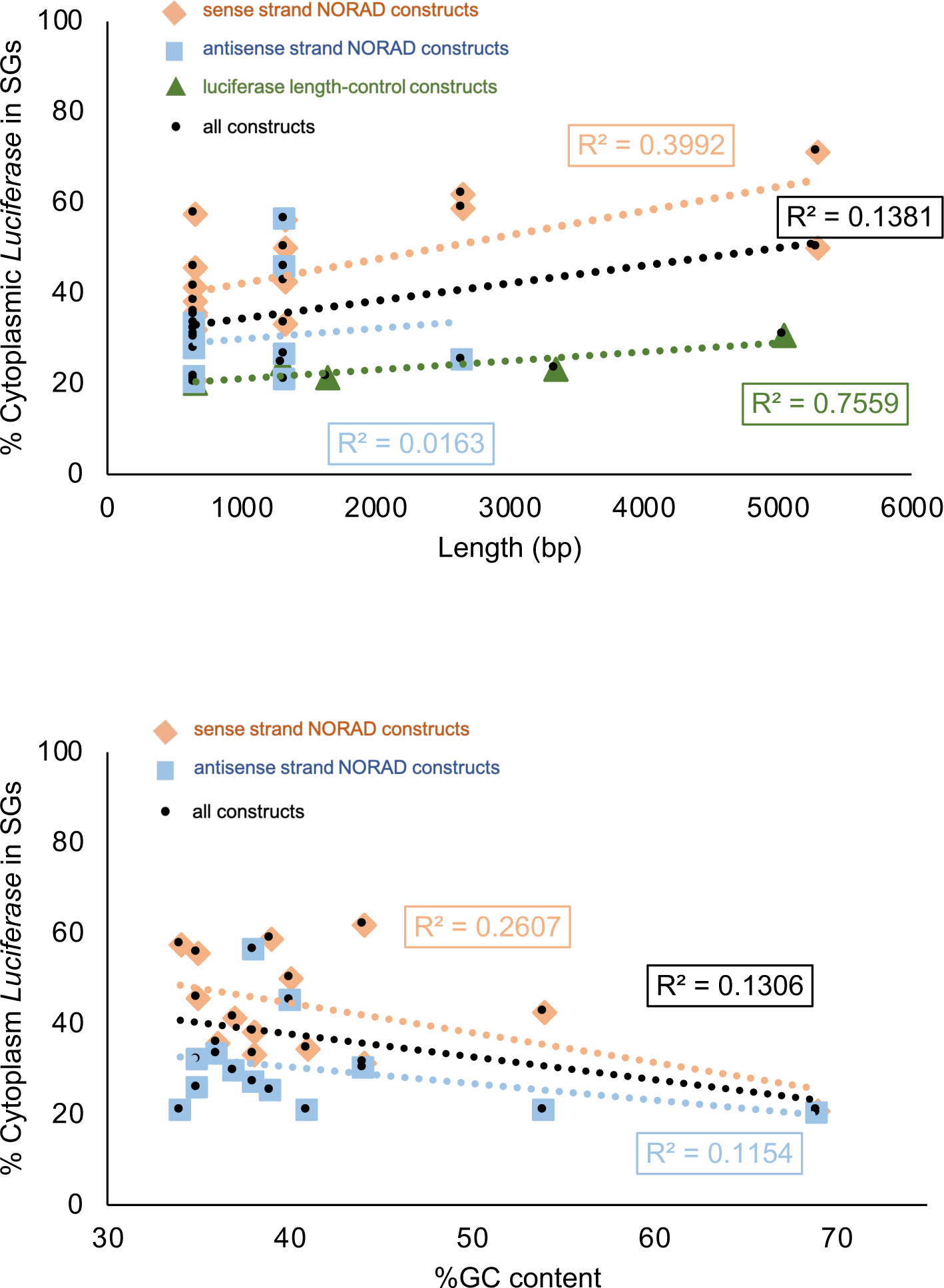
Correlation between SG enrichment and length/GC content **(A)** Correlation between SG enrichment and constructs’ length. Orange diamonds are sense-NORAD constructs, light blue squares are antisense-NORAD constructs, dark green triangles are luciferase length control constructs, and black circles are all the constructs. (**B**) Correlation between SG enrichment and %GC content of the contructs. Orange diamonds are sense-NORAD constructs, light blue squares are antisense-NORAD constructs, and black circles are all the constructs.

**Figure S3:**
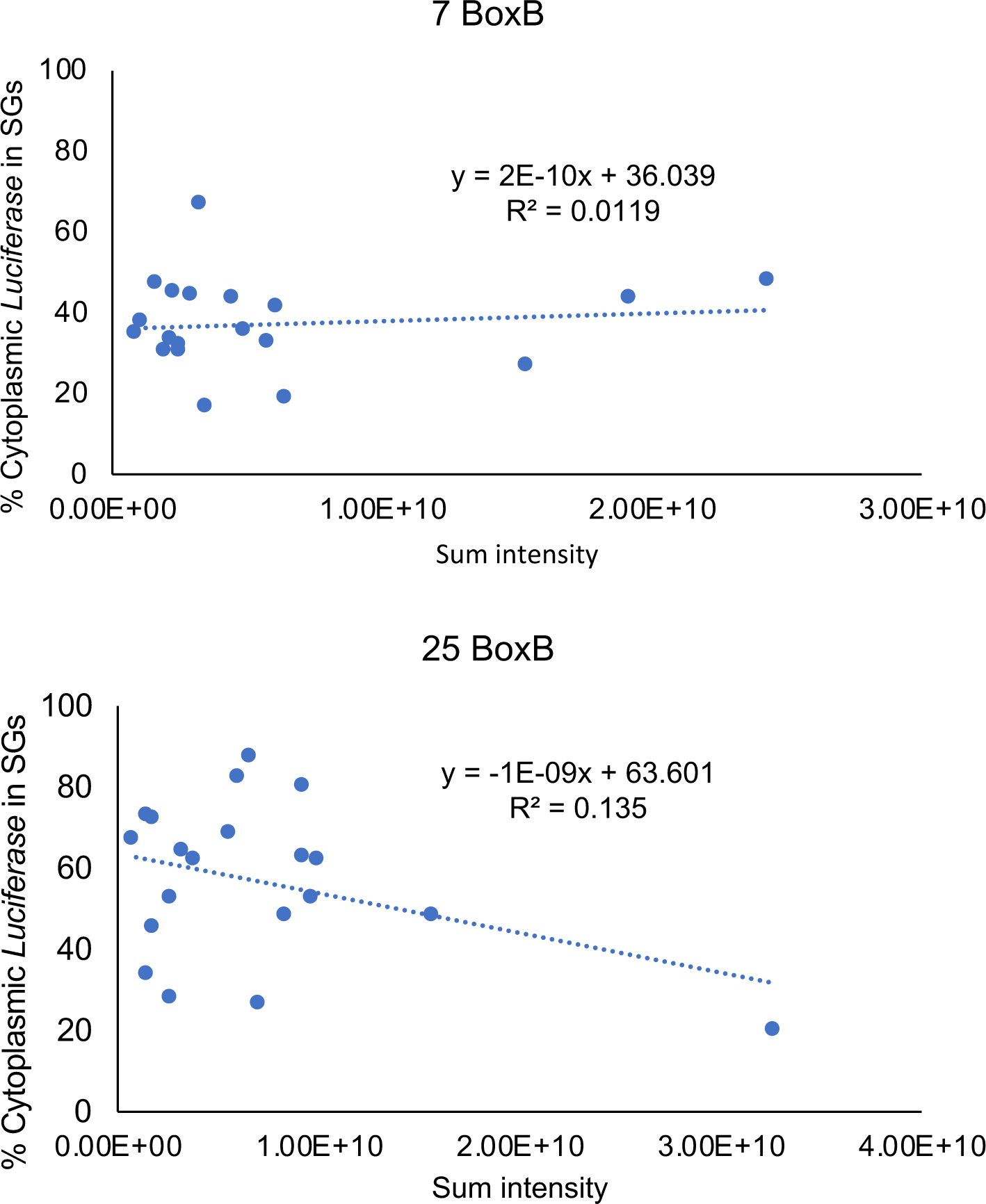
Correlation between G3BP intensity and *luciferase* SG enrichment **(A)** Correlation between SG enrichment of *luciferase* with 7 BoxB and expression level of G3BP1-GFP-λN. (**B**) Correlation between SG enrichment of *luciferase* with 25 BoxB and expression level of G3BP1-GFP-λN.

**Figure S4:**
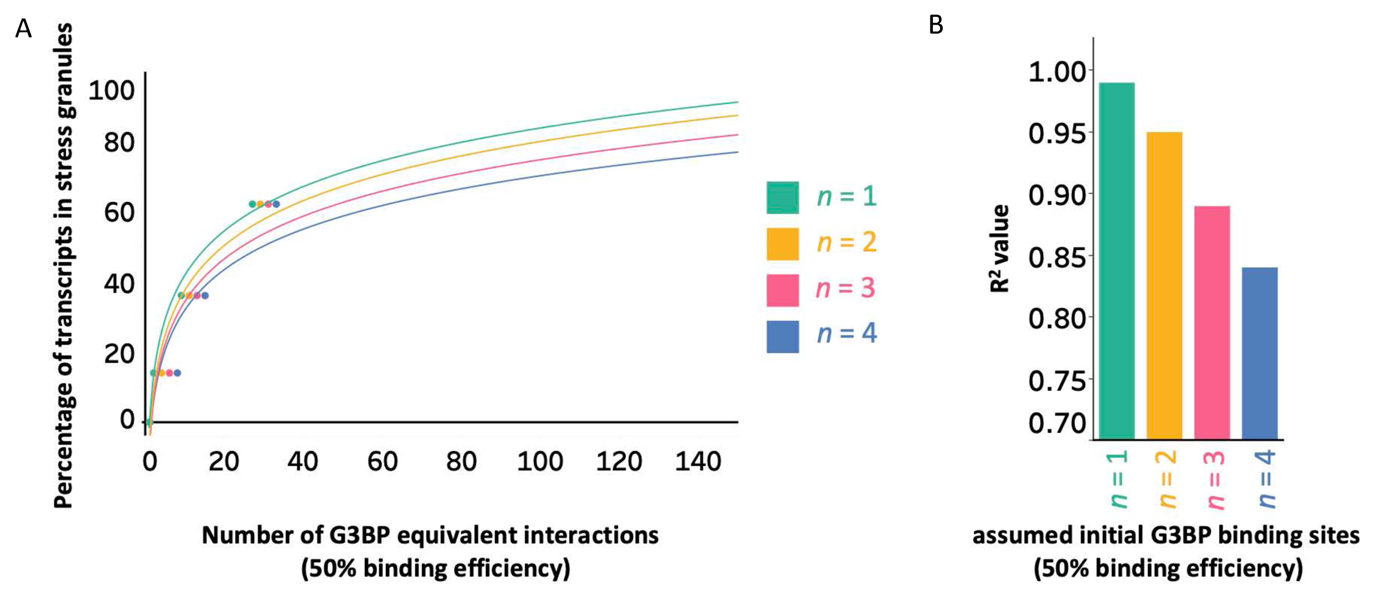
Mathematical modeling of RBP interactions assuming 50% occupancy **(A)** Curve fitting of percentage of *luciferase* in SGs vs. number of G3BP tethering sites, assuming only half of the tethering sites are occupied. **(B)** Bar graph showing how r^2^ changes with n.

**Figure S5:**
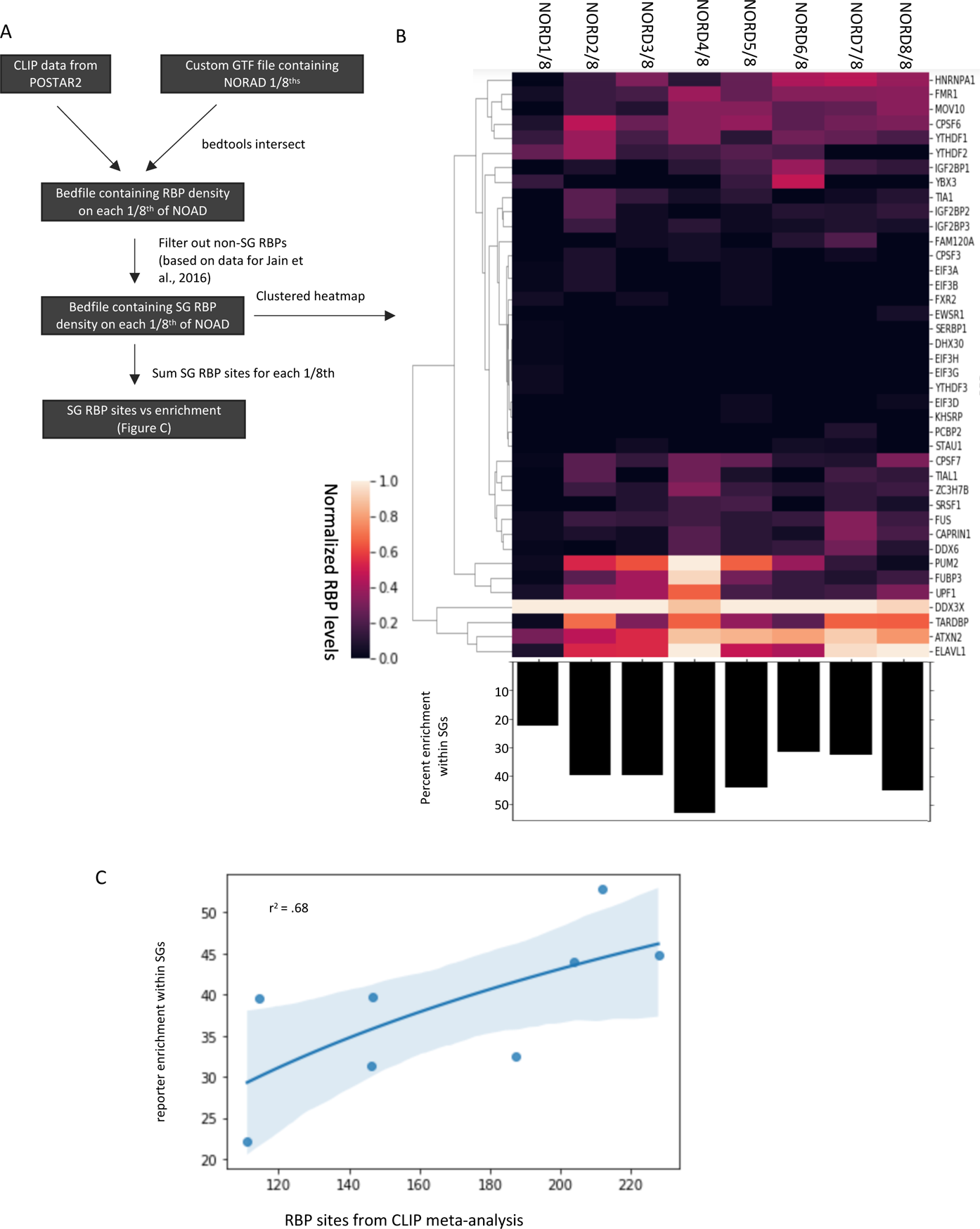
CLIP analysis of NORAD RBP interactions **(A)** Schematic depicting how RBP CLIP analysis was performed. **(B)** *Top:* Clustered heatmap depicting RBP CLIP sites for each 1/8^th^ of the NORAD transcript. *Bottom:* Barplots showing percentage of transcripts enriched in stress granules. **(C)** Scatterplot showing SG enrichment vs. number of summed SG CLIP sites.

**Figure S6:**
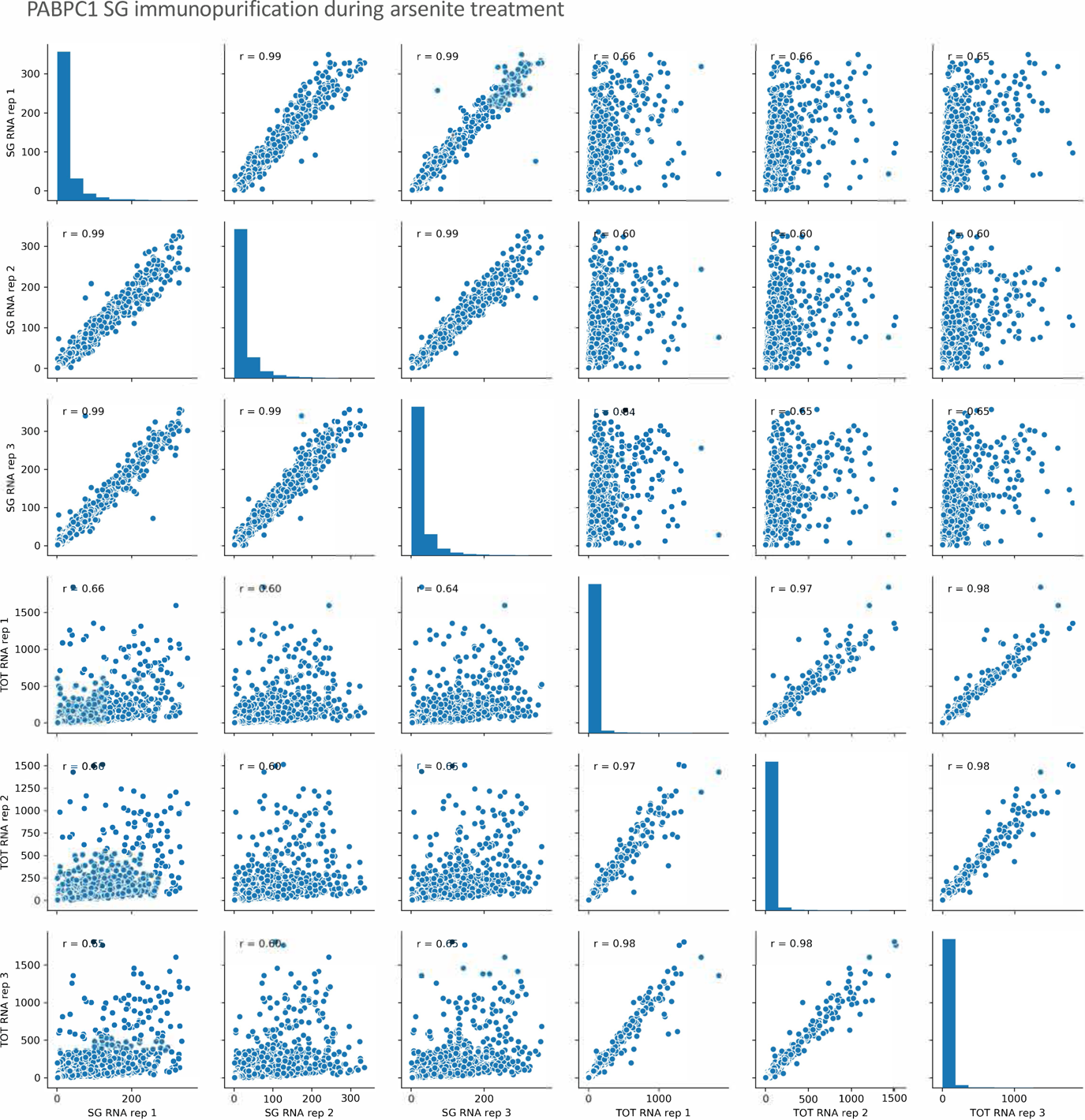
Total RNA and SG core purification via PABPC1 pulldown under arsenite stress yields reproducible transcriptomes Pairwise scatterplots and Pearson correlations for PABPC1 SG immunopurification and total RNA replicates during arsenite treatment.

**Figure S7:**
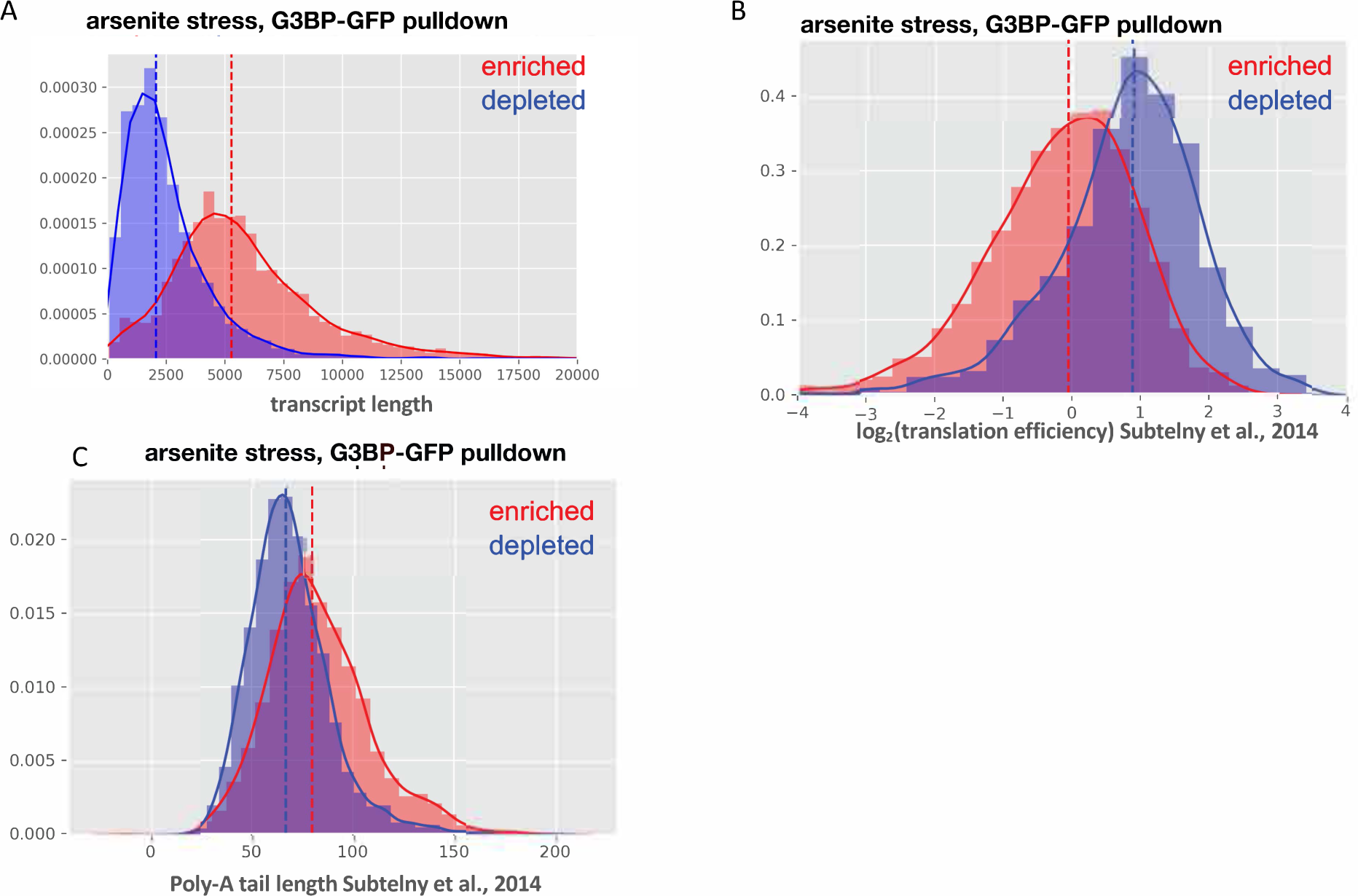
SG cores purified with G3BP immunopurification are enriched for longer RNAs with decreased translation efficiency scores and longer poly-A tails Histograms of (A) transcript length, (B) translation efficiency, and (C) poly-A tail length of SG enriched and depleted transcripts from G3BP1-GFP immunopurification of arsenite-induced SGs (Khong et al., 2017).

**Figure S8:**
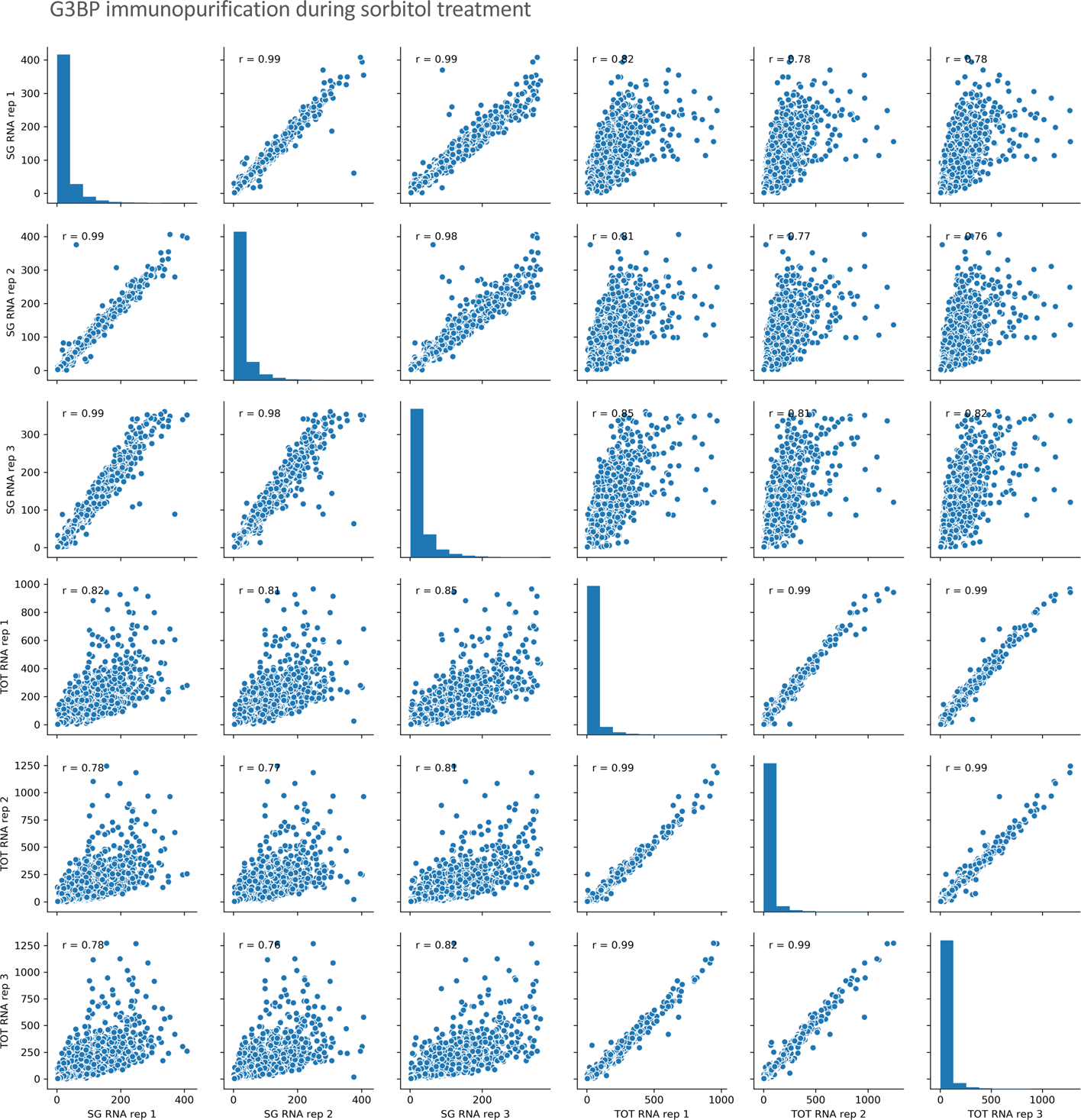
Total RNA and SG core purification via G3BP1-GFP pulldown under sorbitol stress yields reproducible transcriptomes Pairwise scatterplots and Pearson correlations for G3BP SG immunopurification and total RNA replicates during sorbitol treatment.

**Figure S9:**
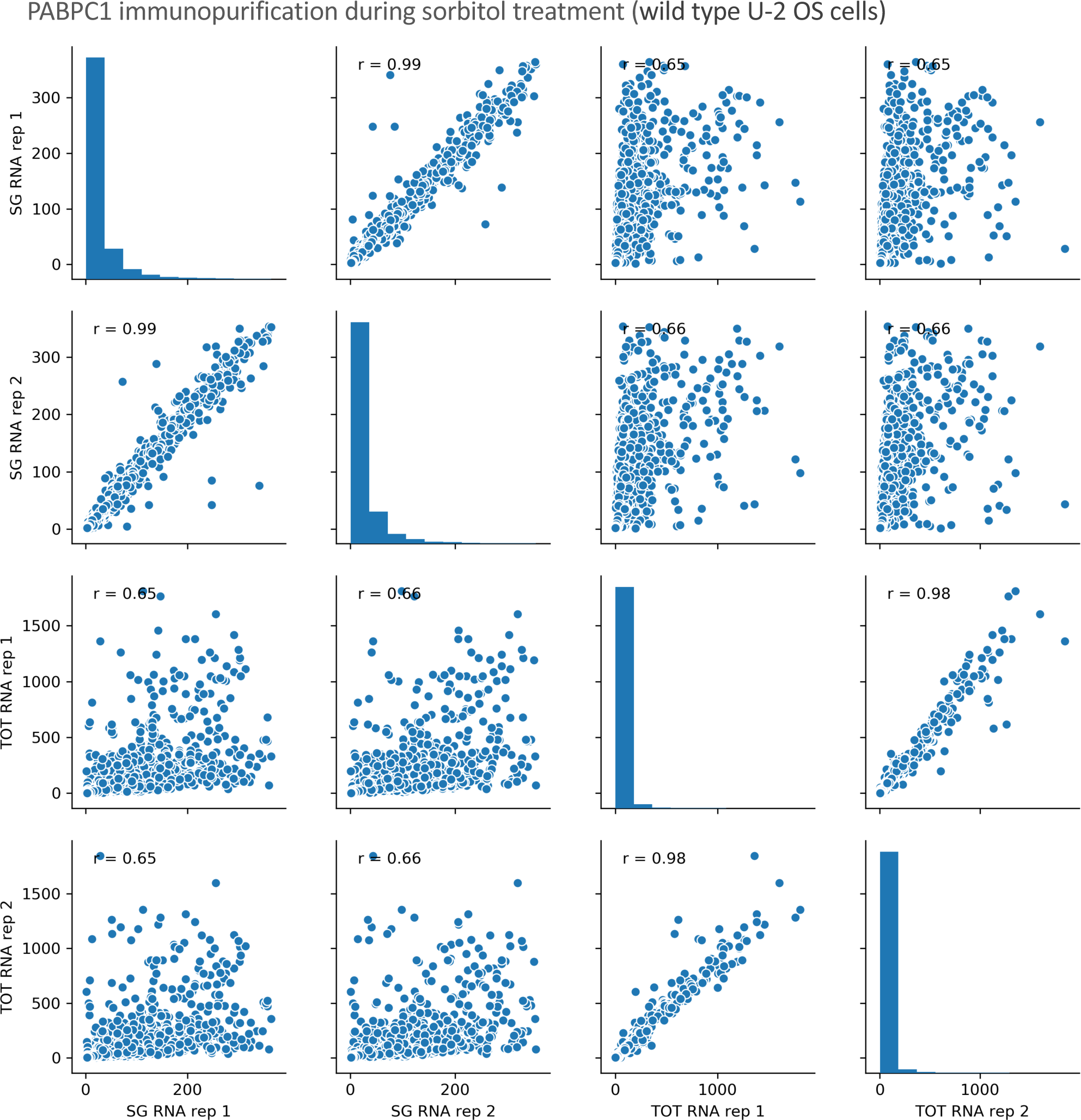
Total RNA and SG core purification via PABPC1 pulldown under sorbitol stress yields reproducible transcriptomes Pairwise scatterplots and Pearson correlations for PABPC1 SG immunopurification and total RNA replicates during sorbitol treatment.

**Figure S10:**
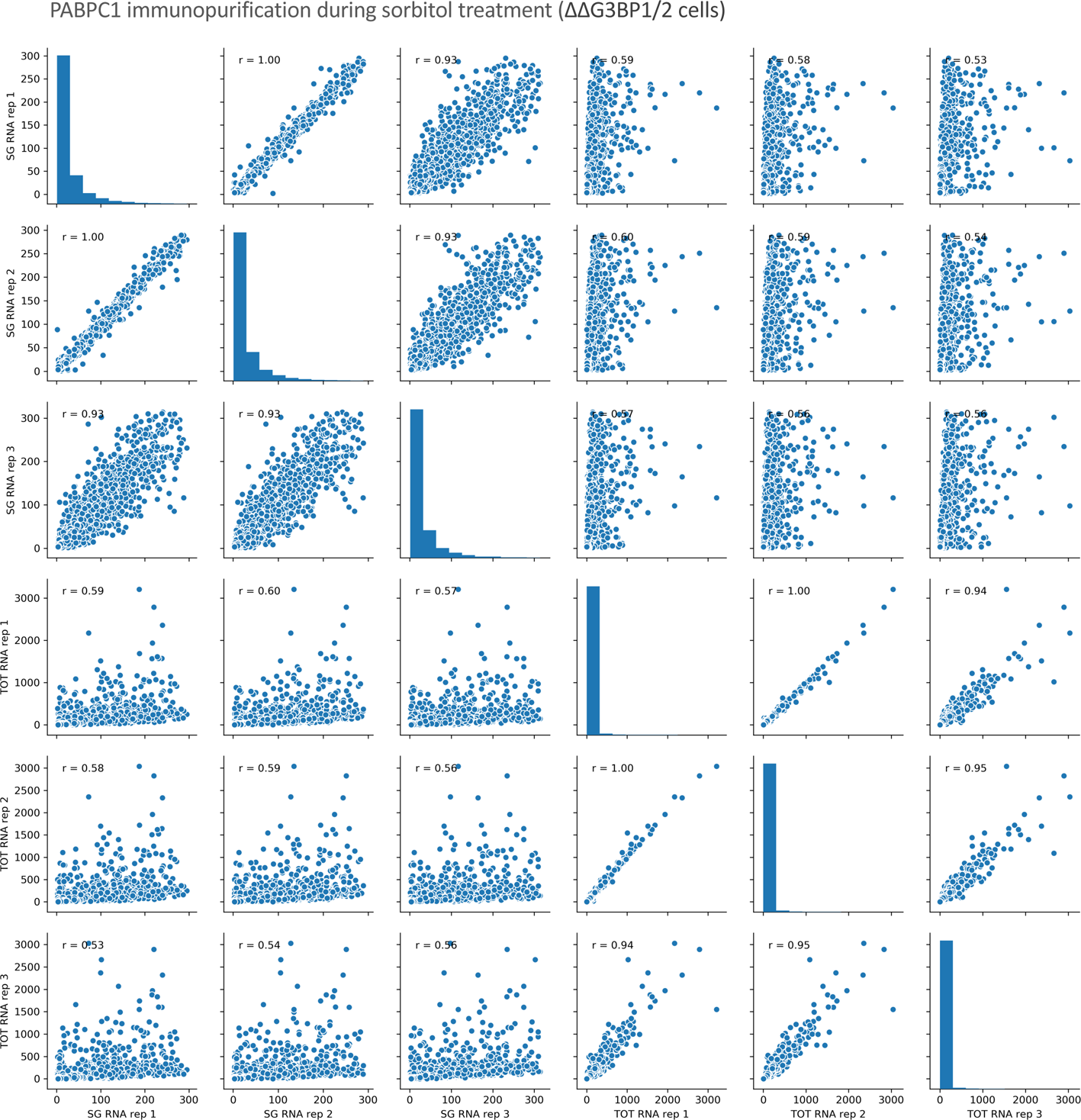
Total RNA and SG core purification from ΔΔG3BP1/2 cells via PABPC1 pulldown under sorbitol stress yields reproducible transcriptomes Pairwise scatterplots and Pearson correlations for PABPC1 SG immunopurification and total RNA replicates during sorbitol treatment in ΔΔG3BP1/2 cells.

**Figure S11:**
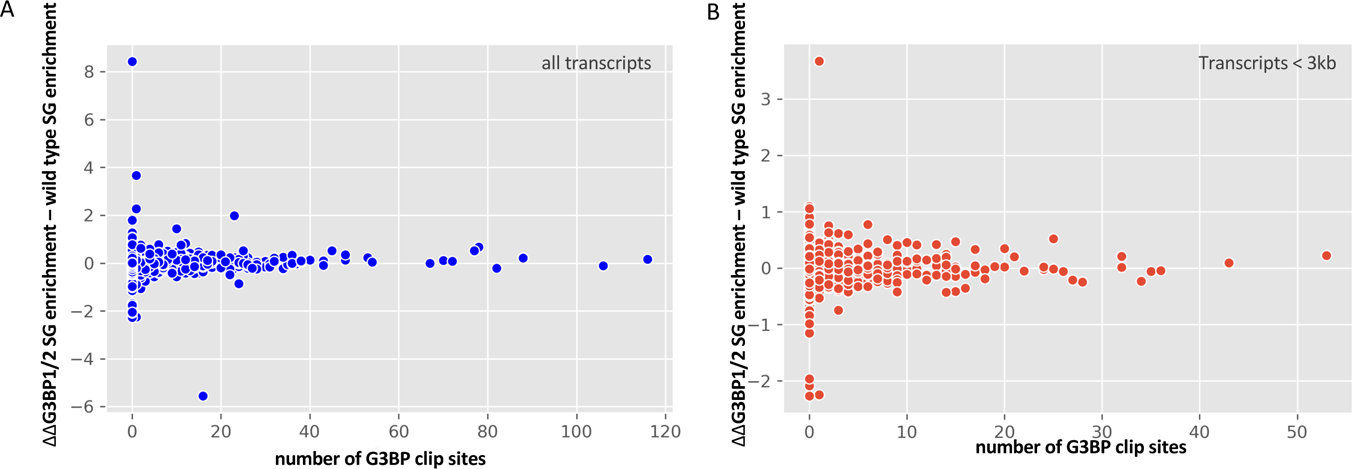
Analysis of the change in SG enrichment scores in G3BP deletion cells as a function of the number of G3BP CLIP sites. **(A)** ΔΔG3BP1/2 SG enrichment – wild type SG enrichment vs. number of G3BP CLIP sites. **(B)** Same as A, but only for transcripts < 3 kb.

**Figure S12:**
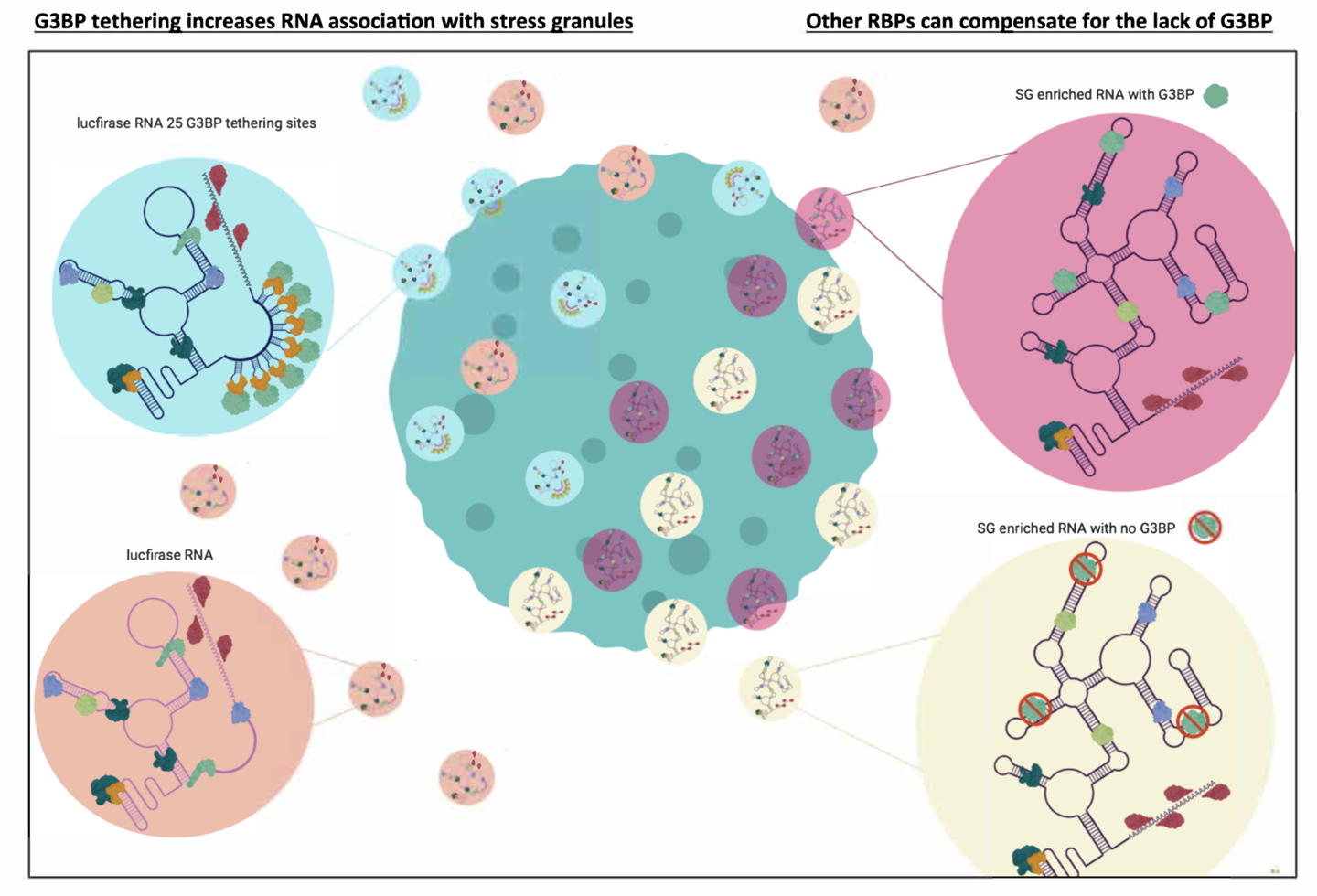
Synergistic recruitment of mRNPs to SGs based on the number of SG interactions Model for RNA recruitment for stress granules. ***Left:*** Tethering, e.g. G3BP to RNAs that contain few SG interactions, can increase an RNA’s association with stress granules. ***Right:*** Deletion of individual interactions from RNPs with many interactions with stress granules leads to limited changes in RNA recruitment to SGs since other interactions can compensate.

**Table S1:** Table of all luciferase-NORAD chimera constructs. This table lists all constructs used in the NORAD experiments with length, SG enrichment, and predicted protein binding sites (assuming luciferase has 2 protein binding sites).

| Construct | Length (bp) | %SG accumulation | Predicted number of protein binding sites |
| --- | --- | --- | --- |
| FL NORAD | 5287 | 70.9 | 51 |
| NORAD-PREmut | 5287 | 50 | 33 (assuming SAM68 sites not affected) |
| 1/2 NORAD | 2644 | 61.8 | 24 |
| 2/2 NORAD | 2643 | 58.9 | 29 |
| 1/4 NORAD | 1322 | 42.6 | 9 |
| 2/4 NORAD | 1322 | 56 | 17 |
| 3/4 NORAD | 1322 | 50 | 22 |
| 4/4 NORAD | 1321 | 33.2 | 9 |
| 1/8 NORAD | 661 | 20.9 | 2 |
| 2/8 NORAD | 661 | 38.2 | 9 |
| 3/8 NORAD | 661 | 35.8 | 7 |
| 4/8 NORAD | 661 | 57.5 | 12 |
| 5/8 NORAD | 661 | 41.3 | 15 |
| 6/8 NORAD | 661 | 31.8 | 8 |
| 7/8 NORAD | 661 | 34.8 | 6 |
| 8/8 NORAD | 660 | 45.6 | 5 |
| 2/2 asNORAD | 2643 | 25 | 8 |
| 1/4 asNORAD | 1322 | 20.7 | 4 |
| 2/4 asNORAD | 1322 | 26.2 | 5 |
| 3/4 asNORAD | 1322 | 45.4 | 3 |
| 4/4 asNORAD | 1321 | 56.25 | 7 |
| 1/8 asNORAD | 661 | 20 | 2 |
| 2/8 asNORAD | 661 | 27.45 | 4 |
| 3/8 asNORAD | 661 | 33.3 | 4 |
| 4/8 asNORAD | 661 | 21.2 | 3 |
| 5/8 asNORAD | 661 | 29.8 | 3 |
| 6/8 asNORAD | 661 | 30.4 | 2 |
| 7/8 asNORAD | 661 | 21.1 | 4 |
| 8/8 asNORAD | 660 | 32 | 5 |
| 660bp Luc | 660 | 20.2 | 3 |
| 1300 bp Luc | 1300 | 24.2 | 4 |
| FL Luc | 1635 | 21.1 | 4 |
| 2xLuc | 3340 | 23.25 | 6 |
| 3xLuc | 5045 | 30.5 | 8 |

\*Assuming Luciferase has 2 protein binding sites

**Supplemental Table S2: Table of predicted number of RBP sites for all transcripts** This table results from the curve fitting performed in Figure 4 and globally estimates how many ‘G3BP equivalent interactions’ each transcript has with SGs.

**Supplemental Table S3:** Plasmids and Oligos This table describes all plasmids used in this study, including their construction. Oligos used for plasmid construction are also listed.

## Notes

https://www.ncbi.nlm.nih.gov/geo/query/acc.cgi?acc=GSE119977

### Competing Interest Statement

The authors have declared no competing interest.

## REFERENCES

1. Anderson, P., Kedersha, N., and Ivanov, P. (2015). Stress granules, P-bodies and cancer. Biochim. Biophys. Acta-Gene Regul. Mech. 1849, 861–870.

2. Ascano, M. Jr., Mukherjee, N., Bandaru, P., Miller, J.B., Nusbaum, J.D., Corcoran, D.L., Langlois, C., Munschauer, M., Dewell, S., Hafner, M., et al. (2012). FMRP targets distinct mRNA sequence elements to regulate protein expression. Nature 492, 382–386.

3. Baron-Benhamou, J., Gehring, N.H., Kulozik, A.E., and Hentze, M.W. (2004). Using the lambdaN peptide to tether proteins to RNAs. Methods Mol. Biol. 257, 135–154.

4. Buchan, J.R., and Parker, R. (2009). Eukaryotic Stress Granules: The Ins and Outs of Translation. Mol. Cell 36, 932–941.

5. Cirillo, L., Cieren, A., Barbieri, S., Khong, A., Schwager, F., Parker, R., and Gotta, M. (in press). UBAP2L forms distinct cores that act in nucleating stress granules upstream of G3BP1. Current Biology

6. Dewey, C.M., Cenik, B., Sephton, C.F., Dries, D.R., Iii, P.M., Good, S.K., Johnson, B.A., Herz, J., and Yu, G. (2011). TDP-43 Is Directed to Stress Granules by Sorbitol, a Novel Physiological Osmotic and Oxidative Stressor. Mol Cell Biol. 31, 1098–1108.

7. Dunagin, M.C., Cabili, M.N., Rinn, J.L. and Raj, A. (2015). Visualization of lncRNA by Single-Molecule Fluorescence In Situ Hybridization. In Nuclear bodies and Noncoding RNAs: Methods and Protocols, S. Nakagawa and T. Hirose, eds. (New York: Springer Science), pp. 3–19.

8. Gaspar, I., Wippich, F., and Ephrussi, A. (2017). Enzymatic production of single-molecule FISH and RNA capture probes. RNA 23, 1582–1591.

9. Gilks, N., Kedersha, N., Ayodele, M., Shen, L., Stoecklin, G., Dember, L. M., and Anderson, P. (2004). Stress granule assembly is mediated by prion-like aggregation of TIA-1. Molecular Biology of the Cell 15, 5383–5398.

10. Itzhak, D.N., Tyanova, S., Cox, J., and Borner, G.H. (2016). Global, quantitative and dynamic mapping of protein subcellular localization. eLife 5, https://doi.org/10.7554/eLife.16950.001

11. Jain, S., Wheeler, J.R., Walters, R.W., Agrawal, A., Barsic, A., and Parker, R. (2016). ATPase-Modulated Stress Granules Contain a Diverse Proteome and Substructure. Cell 164, 487–498.

12. Kedersha, N., Ivanov, P., and Anderson, P. (2013). Stress granules and cell signaling: more than just a passing phase? Trends Biochem. Sci. 38, 494–506.

13. Kedersha, N., Panas, M.D., Achorn, C.A., Lyons, S., Tisdale, S., Hickman, T., Thomas, M., Lieberman, J., McInerney, G.M., Ivanov, P., et al. (2016). G3BP-Caprin1-USP10 complexes mediate stress granule condensation and associate with 40S subunits. J. Cell Biol. 212, 845–860.

14. Khong, A., Matheny, T., Jain, S., Mitchell, S.F., Wheeler, J.R., and Parker, R. (2017). The Stress Granule Transcriptome Reveals Principles of mRNA Accumulation in Stress Granules. Mol. Cell 68, 808–820.

15. Khong, A., Jain, S., Matheny, T., Wheeler, J.R., and Parker, R. (2018). Isolation of mammalian stress granule cores for RNA-Seq analysis. Methods 137, 49–54.

16. Khong and Parker. (2020). The landscape of eukaryotic mRNPs. RNA 26, 229–239.

17. Kim, H.J., Kim, N.C., Wang, Y., Scarborough, E.A., Moore, J., Diaz, Z., Maclea, K.S., Freibaum, B., Li, S., Molliex, A., et al. (2013). Mutations in prion-like domains in hnRNPA2B1 and hnRNPA1 cause multisystem proteinopathy and ALS. Nature 495, 467– 473.

18. Lee, S., Kopp, F., Chang, T., Yu, H., Xie, Y., and Mendell, J.T. (2016). Noncoding RNA NORAD Regulates Genomic Stability by Sequestering PUMILIO Proteins. Cell 164, 69– 80.

19. Li, Y.R., King, O.D., Shorter, J., and Gitler, A.D. (2013). Stress granules as crucibles of ALS pathogenesis. J. Cell Biol. 201, 361–372.

20. Lima, S.A., Chipman, L.B., Nicholshon, A.L., Chen, Y.H., Yee, B.A., Yeo, G.W., Coller, J., and Pasquinelli, (2017). Short poly(A) tails are a conserved feature of highly expressed genes. Nat. Struct. Mol. Biol. 24, 1057–1063.

21. Markmiller, S., Soltanieh, S., Server, K.L., Mak, R., Jin, W., Fang, M.Y., Luo, E.-C., Krach, R., Yang, D., Sen, A., et al. (2018). Context-Dependent and Disease-Specific Diversity in Protein Interactions with Stress Granules. Cell 172, 590–604.

22. Martin, R.M., Rino, J., Carvalho, C., Kirchhausen, T., and Carmo-Fonseca, M. (2013). Live-cell visualization of pre-mRNA splicing with single-molecule sensitivity. Cell Rep. 4, 1144–1155.

23. Moon, S.L., Morisaki, T., Khong, A., Lyon, K., Parker, R., and Stasevich, T.J. (2019). Multicolour single-molecule tracking of mRNA interactions with RNP granules. Nature Cell Biology 21, 162–168.

24. Namkoong, S., Ho, A., Woo, Y.M., Kwak, H., and Lee, J.H. (2018). Systematic Characterization of Stress-Induced RNA Granulation. Mol. Cell 70, 175–187.

25. Niewidok, B., Igaev, M., Pereira, A., Strassner, A., Lenzen, C., Richter, C.P., and Piehler, J. (2018). Single-molecule imaging reveals dynamic biphasic partition of RNA-binding proteins in stress granules. JBC 217, 1303–1318.

26. Nihongaki, Y., Yamamoto, S., Kawano, F., Suzuki, H., and Sato, M. (2015). CRISPR-Cas9-based Photoactivatable Transcription System. Chem Biol. 22, 169–174.

27. Panas, M.D., Ivanov, P., and Anderson P. (2016). Mechanistic insights into mammalian stress granule dynamics. J. Cell Biol. 7, 313–323.

28. Protter, D.S.W., and Parker, R. (2016). Principles and Properties of Stress Granules. Trends Cell Biol. 26, 668–679.

29. Reineke, L.C., and Lloyd, R.E. (2013). Diversion of stress granules and P-bodies during viral infection. Virology 436, 255–267.

30. Schindelin, J., Arganda-Carreras, I., Frise, E., Kaynig, V., Longair, M., Pietzsch, T., Preibisch, S., Rueden, C., Saalfeld, S., Schmid, B., et al. (2012). Fiji: an open-source platform for biological-image analysis. Nat Methods 9, 676-682.

31. Somasekharan, S.P., El-Naggar, A., Leprivier, G., Cheng, H., Hajee, S., Grunewald, T.G.P., Zhang, F., Ng, T., Delattre, O., Evdokimova, V., et al. (2015). YB-1 regulates stress granule formation and tumor progression by translationally activating G3BP1. J. Cell Biol. 208, 913–929.

32. Subtelny, A.O., Eichhorn, S.W., Chen, G.R., Sive, H., and Bartel, D.P. (2014). Poly(A)-tail profiling reveals an embryonic switch in translational control. Nature 508, 66–71.

33. Tauber, D., Tauber, G., Khong, A., Van Treeck, B., Pelletier, J., and Parker, R. (2020). Modulation of RNA condensation by the DEAD-box protein eIF4A. Cell 180, 411–426.

34. Tichon, A., Gil, N., Lubelsky, Y., Havkin Solomon, T., Lemze, D., Itzkovitz, S., Stern-Ginossar, N., and Ulitsky, I. (2016). A conserved abundant cytoplasmic long noncoding RNA modulates repression by Pumilio proteins in human cells. Nat Commun. 7, doi:10.1038/ncomms12209.

35. Tichon, A., Perry, R. B.-T., Stojic, L., and Ulitsky, I. (2018). SAM68 is required for regulation of Pumilio by the NORAD long noncoding RNA. Genes & Dev. 32, 70–78.

36. Tourrière, H., Chebli, K., Zekri, L., Courselaud, B., Blanchard, J.M., Bertrand, E., and Tazi, J. (2003). The RasGAP-associated endoribonuclease G3BP assembles stress granules. J. Cell Biol. 160, 823–831.

37. Trapnell, C., Hendrickson, D.G., Sauvageau, M., Goff, L., Rinn, J.L., and Pachter, L. (2013). Differential analysis of gene regulation at transcript resolution with RNA-seq. Nat Biotech 31, 46–53.

38. Van Nostrand, E.L., Freese, P., Pratt, G.A., Wang, X., Wei, X., Xiao, R., Blue, S.M., Chen, J.-Y., Cody, N.A.L., Dominguez, D., et al. A Large-Scale Binding and Functional Map of Human RNA Binding Proteins. BioRxiv 179648.

39. Van Treeck, B., Protter, D.S.W., Matheny, T., Khong, A., Link, C.D., and Parker, R. (2018). RNA self-assembly contributes to stress granule formation and defining the stress granule transcriptome. PNAS 115, 2734–2739.

40. Wheeler, J.R., Matheny, T., Jain, S., Abrisch, R., and Parker, R. (2016). Distinct stages in stress granule assembly and disassembly. eLife 5, https://doi.org/10.7554/eLife.18413.001.

41. Wheeler, J.R., Jain, S., Khong, A., and Parker, R. (2017). Isolation of yeast and mammalian stress granule cores. Methods 126, 12–17.

42. Yang, C. Di, B. Hu, M. Zhou, Y. Liu, N. Song, Y. Li, J. Umetsu, Z.J. (2015). LuCLIPdb: a CLIP-seq database for protein-RNA interactions BMC Genomics 16, 51.

43. Youn, J., Dunham, W.H., Hong, S.J., Fabian, M., Youn, J., Dunham, W.H., Hong, S.J., Knight, J.D.R., Bashkurov, M., and Chen, G.I. (2018). High-Density Proximity Mapping Reveals the Subcellular Organization of mRNA-Associated Granules and Bodies. Mol. Cell 69, 517–532.

